# Integrating ethnolinguistic and archaeobotanical data to uncover the origin and dispersal of cultivated sorghum in Africa: a genomic perspective

**DOI:** 10.1101/2025.04.16.648676

**Authors:** Aude Gilabert, Monique Deu, Louis Champion, Philippe Cubry, Armel Donkpegan, Jean-François Rami, David Pot, Yves Vigouroux, Christian Leclerc

## Abstract

Archaeobotanical evidence suggests that the beginning of cultivation and emergence of domesticated sorghum was located in eastern Sudan during the fourth millennium BCE. Here, we used a genomic approach, together with archaeobotanical and ethnolinguistic data, to refine the spatial and temporal origin and the spread of cultivated sorghum in Africa.

We built a probability map of the origin of sorghum domestication in Eastern Africa using genomic data and spatial Bayesian models. The origin was located in Eastern Sudan and Western Ethiopia, in perfect concordance with recent archaeobotanical evidence. Calibrated on archaeological remains, our genomic-based model suggests that the beginning of the expansion of sorghum agriculture took place around 4,600 years ago.

Spread of sorghum cultivation led to a sorghum population structure fitting ethnolinguistic groups at different scales, suggesting that human social groups and sorghum populations co-diffused. Consequently, ethnolinguistic barriers and social preferences, as well as adaptation to specific climate zones, have contributed to structuring domesticated sorghum diversity during its diffusion.

## Introduction

Sorghum is one of the most climate-resilient and drought-tolerant cereals. It is the fifth most widely cultivated cereal globally (FAO, 2023) and is the mainstay for millions of inhabitants in the drier areas of Africa and Asia. Sorghum has a multitude of diversified end uses, including food, feed, fodder, fiber, and fuel in over 100 countries in a variety of environments.

Anthropological evidence suggests that hunter-gatherers were familiar with wild forms of sorghum as early as 8,000 BCE (Smith and Frederiksen, 2000). The beginning of cultivation and emergence of sorghum domesticated characteristics was situated during the fourth millennium BCE in eastern Sudan (Winchell et al., 2017), but no genetic evidence has supported that hypothesis so far, and there are still gaps in our understanding of its early diffusion.

Anthropological and human genetics studies suggest that many cultural and linguistic traits, including agricultural expansion (Diamond and Bellwood, 2003), followed demic diffusion models, i.e. the spread occurred through the movements of people (Hewlett et al., 2002), an hypothesis known for the agricultural diffusion as the “farming/language dispersal hypothesis”. Assuming that the initial sorghum populations were propagated under a demic diffusion model through the movements of people from the crop’s area of origin, and were subsequently maintained through time, it can be hypothesized that sorghum genetic groups should be associated with fine linguistic clusters at their destination.

Many anthropological studies suggest that seed exchanges and seed circulation are related to farmers’ linguistic identity (Leclerc and Coppens d’Eeckenbrugge, 2012; Westengen et al., 2014). As noted by Harlan & Stemler (1976), the correspondence between the distribution of the basic morphological groups of sorghum and the major linguistic groups in Africa may not be fortuitous. For example, guinea sorghum is associated with Western African farmers, durra sorghum is confined to the Muslim and Arabic farmers along the fringes of the Sahara and Ethiopia (Kimber, 2000), and kafir, which is widely grown in Southern Africa, is reputed to be associated with the Bantu-speaking peoples of Eastern and Southern Africa (de Wet and Huckabay, 1967). If such social barriers were at work in the past and maintained, at least partially, up to present times, they can explain why, after thousands of years, sorghum genetic groups still remain associated today with fine linguistic clusters.

The aim of this paper is to shed light on the origin and spread of the cultivated sorghum in Africa by integrating ethnolinguistic, archaeobotanical and genomic data. Based on a multidisciplinary approach and on spatially explicit modeling, we seek to identify the geographic origin of cultivated sorghum, and enlighten its expansion history and dynamics in Africa. The origin and spread of cultivated sorghum were considered to result from both biological and social processes, with two leading questions: firstly, how human social groups could have contributed to the dissemination of initial sorghum populations, and secondly, how linguistic barriers could have subsequently contributed to the isolation of sorghum populations from each other and to the building of its current genetic structure.

To examine the origin and dissemination of African sorghum, we initially investigated the relationships between sorghum populations and ethnolinguistic groups at two classification levels. Subsequently, we assessed the congruence between our genomic-based estimation of the geographic origin of cultivated sorghum and the most recent archaeobotanical findings. By integrating genomic and archaeobotanical data, we finally inferred the timing and rate of the westward and southward expansions from its origin.

## Materials and methods

### Plant material

We used a set of cultivated sorghum accessions [*Sorghum bicolor* (L.) Moench] included in several genotype-by-sequencing (GBS) datasets previously published in Brenton et al. (2016), Lasky et al. (2015) and Yu et al. (2016). From these datasets combining over 2,000 landraces, we selected 210 cultivated varieties (Supplementary Dataset 1 available on the CIRAD Dataverse repository; see the section “Data and code availability”) with the aim of 1) keeping only African geo-referenced accessions, 2) covering the African area of distribution of sorghum and 3) limiting over-representation of any geographic, ethnolinguistic and genetic group. To proceed to that selection, we first excluded the non-African landraces from the complete datasets. Then, following the procedure described in the following section “Genomic dataset”, we produced a VCF file containing the African landraces from the Brenton, Morris-Lasky and Yu collections and performed an individual-based clustering analysis implemented in STRUCTURE (Pritchard et al., 2000) and a Neighbor Joining tree based on that African dataset, to remove nearest neighbor accessions while maximizing geographic distance and ethnolinguistic data (results not shown). Latitude and longitude coordinates associated with each selected accession were checked using passport data and place names from different databases (genesis, available at https://www.genesys-pgr.org/a/overview, grin global available at https://www.grin-global.org/help_pw2/Content/Searching%20Overview.htm, and geonames available at https://www.geonames.org/). This sampling included representation of all five known cultivated sorghum botanical groups of cultivated sorghum (*bicolor, caudatum, durra, guinea, and kafir*) and their intermediate races.

### Genomic dataset

The GBS data of cultivated sorghum from Brenton, Morris-Lasky and Yu collections were downloaded from NCBI Sequence Read Archive (Leinonen et al., 2011) (SRA) (see Supplementary Dataset 1). The SRA files were converted to FASTQ file using fastq-dump of SRA Toolkit (http://www.ncbi.nlm.nih.gov/Traces/sra/sra.cgi?view=toolkit_doc&f=fastq-dump).

Low quality bases, Illumina adapters, primers and the end of read were removed with TRIMMOMATIC version 0.33 (Bolger et al., 2014) with the following options: ILLUMINACLIP 2:30:10, LEADING 3, TRAILING 3, SLIDINGWINDOW 4:15, MINLEN 36.

All single-end reads were aligned to the BTx623 sorghum reference genome (McCormick et al., 2018; release 3.1 of the Phytozome database) with the Burrow-Wheeler Aligner (BWA) software v0.7.15 using the standard MEM option (Li and Durbin, 2009). The unmapped, not properly mapped and duplicated reads were filtered with the SAMtools v0.1.9 software (Li et al., 2009). The resulting BAM files (binary format of sequence data) were used as input for Genome Analysis Toolkit (GATK) v4.1.4.0 (DePristo et al., 2011). The GATK HaplotypeCaller algorithm was used for joint variant discovery and genotyping. Single-sample gVCFs were created using the GATK HaplotypeCaller standard parameters according to GATK Best Practices recommendations (DePristo et al., 2011). Finally, single-sample gVCF files were merged into one VCF file using the GenomicsDBImport tool from GATK followed by GenotypeGVCFs for joint variant calling.

Raw VCF was filtered with VCFtools v0.1.13 (Danecek et al., 2011) to obtain high-quality SNPs according to the following criteria: only bi-allelic sites, missing data < 40%, minQ (sites with Quality value above) > 60. After applying all filters, we obtained a VCF file containing 95,362 SNPs for the 210 accessions analyzed.

### Data analysis

#### Population structure and diversity

To identify population structure, we used three distinct and complementary approaches: (i) a maximum likelihood estimation of individual ancestries approach based on a population genetic model as implemented in the software ADMIXTURE (Alexander et al., 2009) (ii) a principal component analysis (PCA) approach; and (iii) an unweighted neighbour-joining (NJ) approach based on pairwise genetic distances. These analyses were run on the dataset of 210 accessions and 95,362 SNPs. ADMIXTURE v.1.22 software (Alexander et al., 2009) was used to estimate the proportion of ancestry of each individual sample in each of the *K* considered genetic clusters, *K* ranging from 1 to 20. Ten repetitions of the algorithm were made for each *K*. We used the cross-validation procedure implemented in ADMIXTURE (5-fold cross-validation) to choose the value of *K* that best represents our sampling. We selected as the optimal number of genetic clusters the value of *K* that gave the lowest cross-validation errors averaged over the 10 replicates. We performed the PCA using PLINK version 1.9 (Chang et al., 2015; Purcell et al., 2007). Diversity tree was constructed by using the unweighted neighbour-joining (NJ) algorithm (Saitou and Nei, 1987) based on a dissimilarity matrix computed using GBS-div software (available at https://darwin.cirad.fr/gbsdiv.php): dissimilarity indices *d_ij_* were estimated by simple matching, that is the number of unmatching alleles between two accessions standardized by the number of loci (Perrier and Jacquemoud-Collet, 2019).

We estimated admixture graphs of geographically defined genetic groups of cultivated sorghum resulting from Admixture analysis using TreeMix software v1.13 (Pickrell and Pritchard, 2012) TreeMix builds a maximum likelihood tree for the relationships among populations and allows the addition of migration events to improve model fit. It allows inference of contemporary and historical patterns of gene flow among populations. We used the script provided by Fitak (2021) to run TreeMix for a number of migration events, *m*. We tested *m* ranging from 0 to 10 with 30 iterations for each value of *m*. We also considered different sizes of SNP-blocks ranging from 50 to 500 with an incrementation of 50, to avoid converging to the same likelihood across all runs for each iteration of *m*, as suggested (Fitak, 2021). The optimal number of migration events was then identified using the Evanno’s method (Evanno et al., 2005) implemented in the R package optM (Fitak, 2021) The tree inferred with the optimal number of migration events was visualized using the R function provided with the Treemix software (Pickrell and Pritchard, 2012). Because of its divergence and its likely independent origin (Mace et al., 2013; Sagnard et al., 2011), we excluded the cluster cl5c for the Treemix analysis.

In order to characterize the level of diversity within cultivated sorghum, we computed the observed heterozygosity (*ho*) and gene diversity (*hs*) per genetic cluster using the R package hierfstat (Goudet, 2005). We calculated the number of singletons and private doubletons per accession using VCFtools 0.1.16 (Danecek et al., 2011) and the number of private alleles (*n*) using an in-house script (available on the CIRAD gitlab platform; see the section “Data and code availability”). The genetic differentiation between the genetic clusters was estimated by the means of pairwise Weir & Cockerham *fst* using the R package hierfstat (Goudet, 2005). All those statistics were computed using the whole dataset of 210 accessions. For all the subsequent analyses that are based on the genetic clusters, we only kept accessions that were unambiguously assigned to a genetic cluster using the same ADMIXTURE ancestry threshold of 0.7.

To analyze the distribution of the genetic clusters according to the climate and to the language families, we used the Köppen-Geiger climate classification, which defines climate zones with different vegetation growth according to the temperature and/or the dryness (Beck et al., 2018), and the language classification referenced in the database Ethnologue Version 16 (Lewis, 2009). Regarding the climate, our sampling is distributed in ten of the sixteen zones present in Africa, with six climate zones being represented by more than five accessions. For the ethnolinguistic analysis, we considered the three main language families present in Africa: the Afro-Asiatic one, distributed in Africa in the western and eastern parts of the Sahelian belt and in the Horn of Africa, the Nilo-Saharan language, whose distribution mostly extends from the central part of the Sahara and Sahelian belt to northern limit of the Great Lakes region, and the Niger-Congo language family, widely distributed from the Sahelian belt to South Africa. We further subdivided the Niger-Congo family into two classes, the Mande distributed in the western part of western Africa and the Atlantic-Congo languages, which include the Bantoid languages notably, since this level of classification has been shown to be relevant, at least for the fonio in Western Africa (Diop et al., 2023). We further subdivided the Nilo-Saharan family into three classes (Central and Eastern Sudanic and Fur), the Afro-Asiatic family into four classes (the Chadic, the Cushitic, the Omotic and the Semitic) and the Atlantic-Congo class into 16 classes to consider the Guthrie classification of the Bantoid languages (Guthrie, 1971, 1948; Maho, 2009).

The association between genetic clusters and botanical races, climate zones or ethnolinguistic groups were tested using Fisher’s exact tests with the R function fisher.test. The associations were further analyzed by visualizing the Peason residuals to identify the specific groups contributing to the observed associations. Pearson residuals were retrieved from the R function chisq.test. Cells with residuals greater than |2| were considered as deviating from the expected values. Kruskal-Wallis rank sum tests (using the R function kruskal.test) were used to test whether the altitude or the number of singletons per accession differed between the genetic clusters. Multiple groups comparisons were performed using Dunn’s tests from the R package dunn.test (Dinno, 2024) to identify the genetic clusters that are differing according to the altitude or the number of singletons per accession. All R analyses were performed using R Statistical Software version 4.4.0 (R Core Team, 2024).

#### Inference of the geographic origin of the cultivated sorghum

We used SPLATCHE2 (Currat et al., 2004) to perform spatially-explicit simulations within an approximate Bayesian framework in order to infer the geographic origin of the diffusion of the cultivated sorghum in Africa.

SPLATCHE2 enables users to model range expansions using non-equilibrium stepping-stone models. In those models, the geographical cells are colonized from their neighbors. We considered here a single origin model. We performed 1,000,000 range expansion simulations from randomly picked origins based on prior distributions for latitudinal and longitudinal coordinates, considering a heterogeneous environment and modeling the African continent using an array of 87 by 83 demes. Prior distributions allowed the geographic coordinates of the origin of expansion to vary over the Sub-saharan region, within an area ranging between -17° and 50° of longitude and between -35° and 20° of latitude. Despite the importance of gene flow and introgression in the evolutionary history of sorghum, we decided to consider simple demographic models, to limit the model complexity and the number of parameters in the model. Because domestication can be accompanied by a genetic bottleneck, followed by a period of stable low population size before expansion due to a period of weak cultivation, we considered two different simple demographic models, one with a constant size of population before the expansion and one with a bottleneck before the expansion (*Supplementary Information*, Fig. S1). Uninformative prior distributions were considered for the longitude, latitude, migration rate, the growth rate, the expansion length (Exp_length), the effective population size before expansion (BtSize, corresponding to population size during the bottleneck), the population size before the bottleneck (AncSize) and the duration of the bottleneck in generations (BtDuration) (*Supplementary Information,* Table S1). Simulated genetic variation was surveyed at 210 geographic sites corresponding to the exact sampling locations of cultivated sorghum accessions. A total of 10,000 haploid loci were simulated for each genotype, and an effective mutation rate of 10^−6^ per base pair per generation was considered. Missing data were randomly included in the simulated dataset at the same frequency as the frequency of missing data in the observed dataset (approx. 0.20). The posterior distributions of the geographic and demographic parameters were inferred using an approximate Bayesian computation (ABC) approach relying on a neural network algorithm for posteriors estimation (R package abc (Csilléry et al., 2012)), considering a tolerance rate of 0.5% and implementing 500 neural networks. Parameter estimations were thus performed accepting 5,000 simulations. The summary statistics used for the ABC analysis included a statistic related to the rare variants and the expected heterozygosity in each genetic cluster identified using the ADMIXTURE analysis, as well as eight classes of the Site Frequency Spectrum (SFS), as described in Cubry et al. (2018). The statistic based on the rare variants corresponds to the mean number of singletons per accession within each genetic cluster, relative to the entire dataset. As for the SFS, the eight classes were defined according to the number of times a SNP was detected in the dataset: one time (singleton), two times, three times, four times, five to six times, seven to 12 times, 13 to 28 times and more than 28 times. Two analyses were performed, one considering the 210 cultivated accessions (model including all cultivated genetic clusters) and one without cultivated accessions that likely originated from an independent event and shows signals of introgression with wild sorghum (Gilabert et al., 2023; Mace et al., 2013; Sagnard et al., 2011) (cluster cl5c; see *Supplementary Information*, SI Text section 1.2). We considered for the calculation of the summary statistics only the accessions assigned to a genetic group with an ancestry > 0.7 according to ADMIXTURE analysis. We evaluated the adjustment of our models using the goodness-of-fit approach described in Lemaire et al. (2016) which evaluates the distance between the simulated summary statistics and the observed ones. We used the gfit function of the abc R package (Csilléry et al., 2012) and the mean as the goodness-of-fit statistic to perform the prior predictive checks. We simulated 1,000 additional datasets, sampling the parameters from the posterior distributions obtained from the previous simulations, to compute the 95% credible intervals of the summary statistics and perform the posterior predictive checks.

#### Estimating the timing of the expansion onset

The availability of archaeological data for sorghum allowed us to calibrate temporally our diffusion model following Burgarella et al. (2018) and estimate the timing of the onset of the expansion of the cultivated sorghum in Africa. From the bibliography, we identified African archaeological sites where archaeobotanical remains of cultivated (or pre-domesticated) sorghum could be identified (*Supplementary Information,* Table S2-A), using for each site the oldest estimated dates to get an estimation of the youngest bound for the beginning of the diffusion of cultivated sorghum in Africa. We conducted 5,000 new spatial demographic simulations by randomly sampling the parameters from the posterior distributions estimated in the previous ABC analysis, and monitored the simulated time for the arrival of the expansion wave at each cell where cultivated sorghum were identified by archaeobotanical data. A linear regression was fitted between the observed dates from archaeological sites and the simulated dates at the same geographical coordinates. The intercept of the real time axis was then estimated as the timing of the onset of the expansion. The regression analysis was performed for each simulation, which allowed us to obtain a distribution of the timing of the expansion’s inception. Because domesticated sorghum was suggested to have dispersed through two movements of diffusion (Fuller and Stevens, 2018), a North-South and an East-West one, we performed two independent regression analyses. The first one considered the sites distributed along an Eastern-Western transect from Sudan to Mali (westward diffusion), while the second considered the sites distributed along the North-South axis (southward diffusion, see *Supplementary Information,* Table S2-A). In addition, because the expansion from the origin toward the North may correspond to a different movement of expansion than the one toward the South, and since we only have one archaeological site north of the inferred point of origin, we estimated the onset of the southward expansion removing the archaeological site from Egypt from the latitudinal analysis (see *Supplementary Information*, Table S2-A). Similarly, we also estimated the westward expansion by removing from the longitudinal analysis the archaeological site from Ethiopia, the only site east of the inferred origin (see *Supplementary Information*, Table S2-A).

#### Estimation of the dispersal front speed

We followed Cobo et al. (2019) to estimate the speed of diffusion of the cultivated sorghum in Africa by the means of time-space regressions. We used the same list of archaeological sites considered for the estimation of the beginning of the dispersal of the cultivated sorghum (*Supplementary Information*, Table S2-A). Although based on a low number of archaeological sites, we performed two estimations, for the longitudinal and latitudinal diffusions, similarly to the estimation of the timing of the onset of sorghum expansion. Indeed, one might expect that the front speeds may be different for the two diffusions as the environments and climate do not present the same variability longitudinally and latitudinally.

We fitted two linear regressions between the oldest estimated dates of the archaeological sites and their distance to the estimated point of origin of the expansion of the cultivated sorghum. For the point of origin of the expansion of sorghum, we used the median of the posterior distributions of the longitude and latitude inferred using the ABC approach. Following Cobo et al. (2019), we used the Haversine equation to compute the great-circle distance (the shortest distance between two points on a sphere) between archaeological sites and the point of origin. We estimated the speed as the inverse of the slope of the time-space regression and its standard error (SE) as being equal to the standard error of the slope divided by the slope squared (Cobo et al., 2019; Fort et al., 2004; Jerardino et al., 2014). Because they were based on a low number of sites, we computed for each of the two regressions the 80% confidence-level interval for the speed, similarly to Jerardino et al. (2014): slope ± *t*_1-_*_α_*_/2,_ *_N_*_-2_ * SE_slope_, with *t* being the Student’s *t* value for the confidence level *α* (0.8 here) and *N* the number of sites considered in the regression.

#### Identification of geographic barriers

We estimated effective migration surfaces using the software FEEMS v1.0.0 (Marcus et al., 2021) (available at https://github.com/NovembreLab/feems) to infer geographic barriers and corridors for the cultivated sorghum. FEEMS assumes an isolation-by-distance pattern (IBD), which stipulates that dispersal occurs at a local scale and that two organisms will be more likely genetically close if they are geographically close. FEEMS identifies areas where the genetic distance between two individuals is either higher or lower than expected given the geographic distance separating them and assuming an IBD. FEEMS was run following the pipeline provided with the publication. As for the SPLATCHE analysis, FEEMS was run using the panel of 210 cultivated accessions from which we removed the nine cultivated accessions of the cluster cl5c, as introgression may bias the results.

## Results

### Sorghum population structure and diversity

A panel of 210 georeferenced accessions representative of the African genetic diversity of cultivated sorghum for which GBS data were available was used to study its domestication history as a biological and social process. A total of 95,362 Single Nucleotide Polymorphisms (SNPs) was identified after bioinformatic treatment of the raw genomic data. The genetic structure analysis of population ancestries of our sampling revealed the existence of nine genetic clusters (*Supplementary Information*, SI Text section 1.1, Fig. S2 and S3). The clusters, associated with the known cultivated sorghum botanical races (Fisher’s exact test *p*-value < 0.005; *Supplementary Information*, Fig. S4), were geographically structured (Fig. 1A), with clusters associated with both Köppen-Geiger agroclimatic zones (Beck et al., 2018) and language families (Lewis, 2009) (see below for the detailed results on the association with the agro-climatic zones and language families). One genetic cluster, associated with the caudatum botanical race, was widely distributed (cl1), partially overlapping the other eight genetic clusters. Three genetic clusters were restricted to Western Africa (cl5a, b and c) and were associated with the guinea botanical race. One cluster (cl2b), associated with the kafir race, was restricted to Southern Africa. Two other clusters (cl4 and cl6), respectively associated with the bicolor/durra-bicolor and the caudatum botanical races, were distributed in Eastern Africa. Lastly, two clusters present in Eastern Africa were also found in Southern (cl2a, associated with the guinea race) and Western Africa (cl3, associated with the durra race) (Fig. 1-A, *Supplementary Information*, Fig. S4).

**Figure 1.**
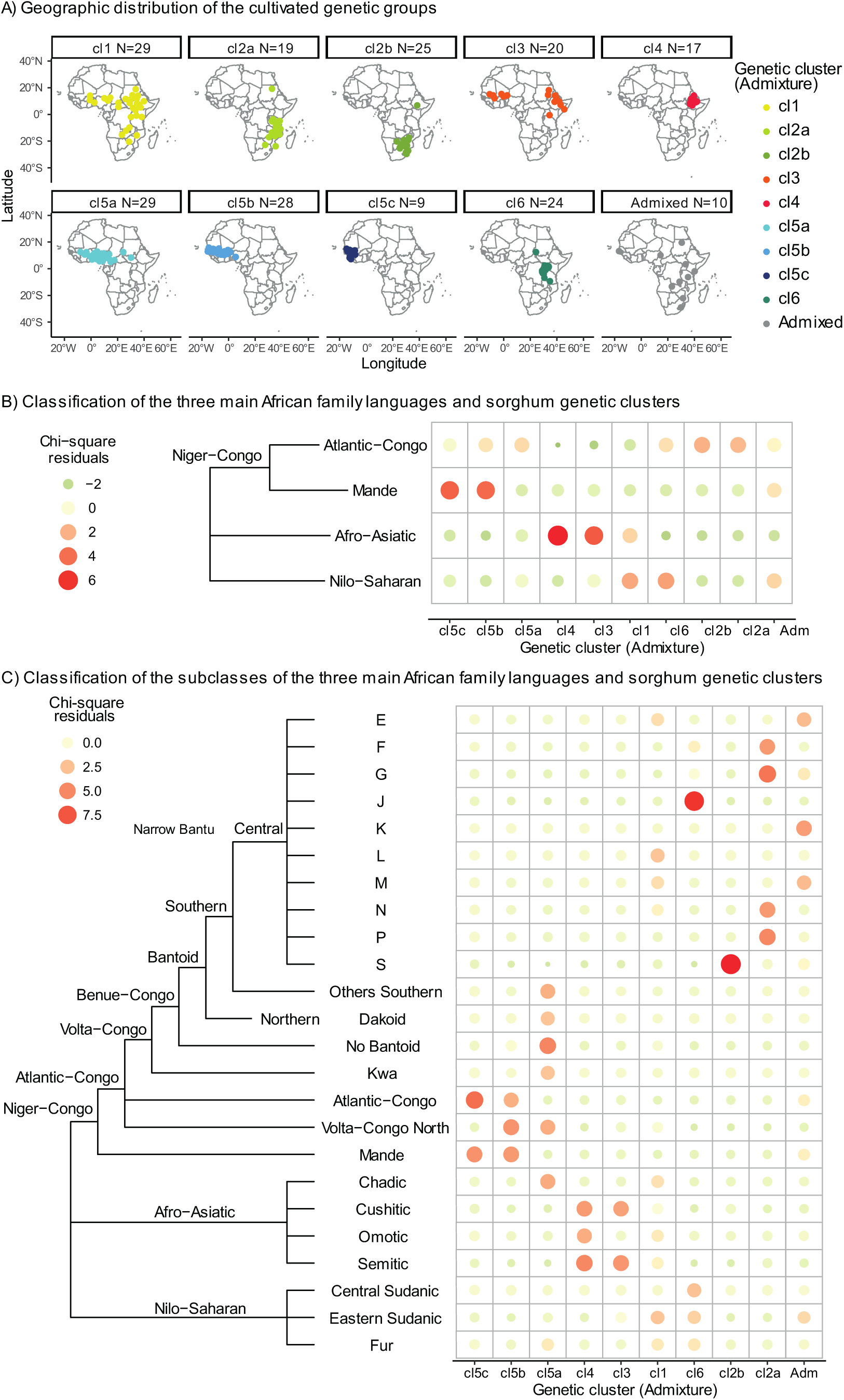
Genetic diversity of the African cultivated sorghum. A) Geographic distribution of the cultivated genetic groups defined using ADMIXTURE (Alexander et al., 2009). Each point corresponds to an accession. B) and C) Chi-squared residuals between the genetic and the farmers language groups, considering (B) four linguistics groups (the three main language families present in Africa, the Niger-Congo family being subdivided into two classes, the Mande and the Atlantic-Congo) or (C) a finer linguistic classification. The size and color of the circles are proportional to the value of the chi-squared residuals between the language families and the genetic clusters.

The genetic clusters considered from the structure analysis (*Supplementary Information*, SI Text section 1.1, Fig. S2 and S3), from the PCA (*Supplementary Information*, SI Text section 1.2, Fig. S5) and from the neighbor joining tree (*Supplementary Information*, Fig. S6) corresponded. The first two axes of the PCA distinguished two distinct groups of accessions that belonged to the Western African cluster cl5c and the Eastern African cluster cl4 (*Supplementary Information*, SI Text section 1.2, Fig. S5-A and C) respectively. A third group, comprising accessions from the Western African clusters cl5a and cl5b, stood apart on the third axis of the PCA (*Supplementary Information*, SI Text section 1.2, Fig. S5-B and C). In the neighbor joining tree (*Supplementary Information*, Fig. S6), the Westernmost cultivated sorghum from the cluster cl5c stood out as a highly divergent and diverse clade, which likely corresponds to accessions belonging to the margaritiferum subrace (see *Supplementary Information*, SI Text section 1.2).

The maximum likelihood tree inferred using the Treemix analysis (Pickrell and Pritchard, 2012) and excluding the highly divergent margaritiferum cultivated cluster cl5c, was consistent with the NJ tree. The analysis suggested two gene flow events, one between the clusters cl1 and cl6 and one between the clusters cl2b and cl3 (*Supplementary Information*, Fig. S7).

Two hotspots of sorghum genetic diversity were localized in Western and Eastern Africa, as illustrated by the distribution of the singletons per accession (*Supplementary Information*, Fig. S8-A). The margaritiferum cluster cl5c in Western Africa harbored the highest levels of genetic diversity compared to all other clusters, as evidenced with the number of singletons per accession (global Kruskal-Wallis rank sum test Χ² = 82.25, df = 9, p < 0.001, Dunn’s tests *p*–values < 0.02 for every pairwise comparison including the cluster cl5c; *Supplementary Information*, Fig. S8-A & B), and the number of private alleles (*Supplementary Information*, Table S3). Besides the cluster cl5c, the Eastern African cluster cl4 and the Eastern and Western African cluster cl3, which harbored similar levels of genetic diversity (Dunn’s test *p*–value for the pairwise comparison cl4-cl3 = 0.41; *Supplementary Information*, Fig. S8-A & B; *Supplementary Information*, Table S3), contributed to the two genetic diversity hotspots.

Sorghum genetic clusters were significantly associated with the Köppen-Geiger agroclimatic zones (Beck et al., 2018) (Fisher’s exact test *p*-value < 0.001; *Supplementary Information*, SI Text section 1.3, Fig. S9-A and B, Table S4-A). The chi-squared residuals of the association between the genetic clusters and the agroclimatic zones indicated that the Eastern African cluster cl4 was overrepresented in temperate climates with warm summers and a dry season (Csb and Cwb zones) and cluster cl3 in hot, arid desert climates (Bwh zone), while Western African clusters cl5c and cl5a were over-represented in the tropical monsoon (Am) and savannah (Aw) climates respectively (*Supplementary Information*, Fig. S9-B, Table S4-A). Conversely, the Eastern African cluster cl2b was under-represented in the tropical savanna climate (Aw) but over-represented in semi-arid (Bsh and Bsk) and Monsoon-influenced humid subtropical climates (Cwa) and cluster cl6 under-represented in hot semi-arid climate (Bsh) but over-represented in humid tropical climate (Af zone) (*Supplementary Information*, Fig. S9-B, Table S4-A). Beside the influence of the agroclimatic zones, the genetic clusters were significantly associated with the elevation (Kruskal-Wallis rank sum test, Kruskal-Wallis Χ² = 120., df = 9, p < 0.001 . The Eastern African cluster cl was found at the highest elevations while the three Western African clusters (cl5b, cl5c) were found at the lowest (*Supplementary Information*, Fig. S9-C).

Association between sorghum genetic clusters and the human language families (Lewis, 2009) was statistically significant (Fisher’s exact test *p*-value < 0.001; Fig. 1-B, *Supplementary Information*, SI Text section 1.3, Fig. S9-D, Table S4-B). The chi-squared residuals showed a clear association between clusters cl2a and cl2b with Atlantic-Congo languages, and between clusters cl5b and cl5c with Mande languages. Clusters cl3 and cl4 were associated with Afro-asiatic languages, and clusters cl1 and cl6 with Nilo-saharan languages (Fig. 1-B). A more precise picture, by considering a finer linguistic classification, revealed that cluster cl2b was strongly (Chi^2^ residual over 9) associated with the Narrow Bantu Guthrie linguistic zone S, which is located in South-Eastern Africa, while the cluster cl2a was associated with four geographically close Bantu Guthrie linguistic zones (zones F, G, N and P). The cluster cl6, still associated with Nilo-saharan languages, was strongly associated with the Narrow Bantu Guthrie linguistic zone J located in the Great Lakes area (*Supplementary Information*, Table S4-B; Fig. 1-C; and map in Fig. S9-D). Finally, the westernmost clusters cl5b and cl5c were associated with the Atlantic-Congo and the Mande, the cluster cl5b being also associated with the Volta-Congo linguistic group. The cluster cl5a was associated with multiple Niger-Congo, non-Bantu, groups and with the Afro-Asiatic Chadic group.

### Geographic origin and spread of the cultivated sorghum

The geographic origin of cultivated sorghum was inferred including or not the cluster cl5c. The cluster cl5c is associated with the margaritiferum subrace (see *Supplementary Information*, SI Text section 1.2), which most probably emerged from a secondary independent domestication event and is likely introgressed with wild sorghum (Gilabert et al., 2023; Mace et al., 2013; Sagnard et al., 2011). Thus, inferring the geographic origin of cultivated sorghum with this cluster could introduce a bias that we wanted to account for (see details in the *Supplementary Information*, SI Text section 2.1 and Fig. S10).

The posterior predictive checks indicate that the model with a constant ancestral population size simulated well and better the observed data than a model with a bottleneck of the ancestral population (*Supplementary Information*, Fig. S10-IA and Fig.S10-IIA). The simplest model with a constant ancestral population size was therefore considered. Graphic representations of the summary statistics (PCA and their distribution) of this model revealed that the posterior distributions were close to the observed dataset, and most of the summary statistics (15 statistics over the 24) were found in the 95% credible intervals of their posterior predictive distributions. Predictive checks are only slightly significant for this no bottleneck model, and more strongly for the other three models (*Supplementary Information*, Table S5). This result was obtained without integrating gene flow between the genetic groups in the models due to methodological limitations of Splatche.

Our retained spatially explicit model (i.e. with constant population effective size before domestication and excluding likely introgressed cluster cl5c) located the geographic origin of cultivated sorghum in a region situated in eastern Sudan and western Ethiopia (Fig. 2-A, *Supplementary Information*, Fig. S10-IB), in line with archaeobotanical evidence and most recent hypotheses (Fuller and Stevens, 2018; Winchell et al., 2017).

**Figure 2.**
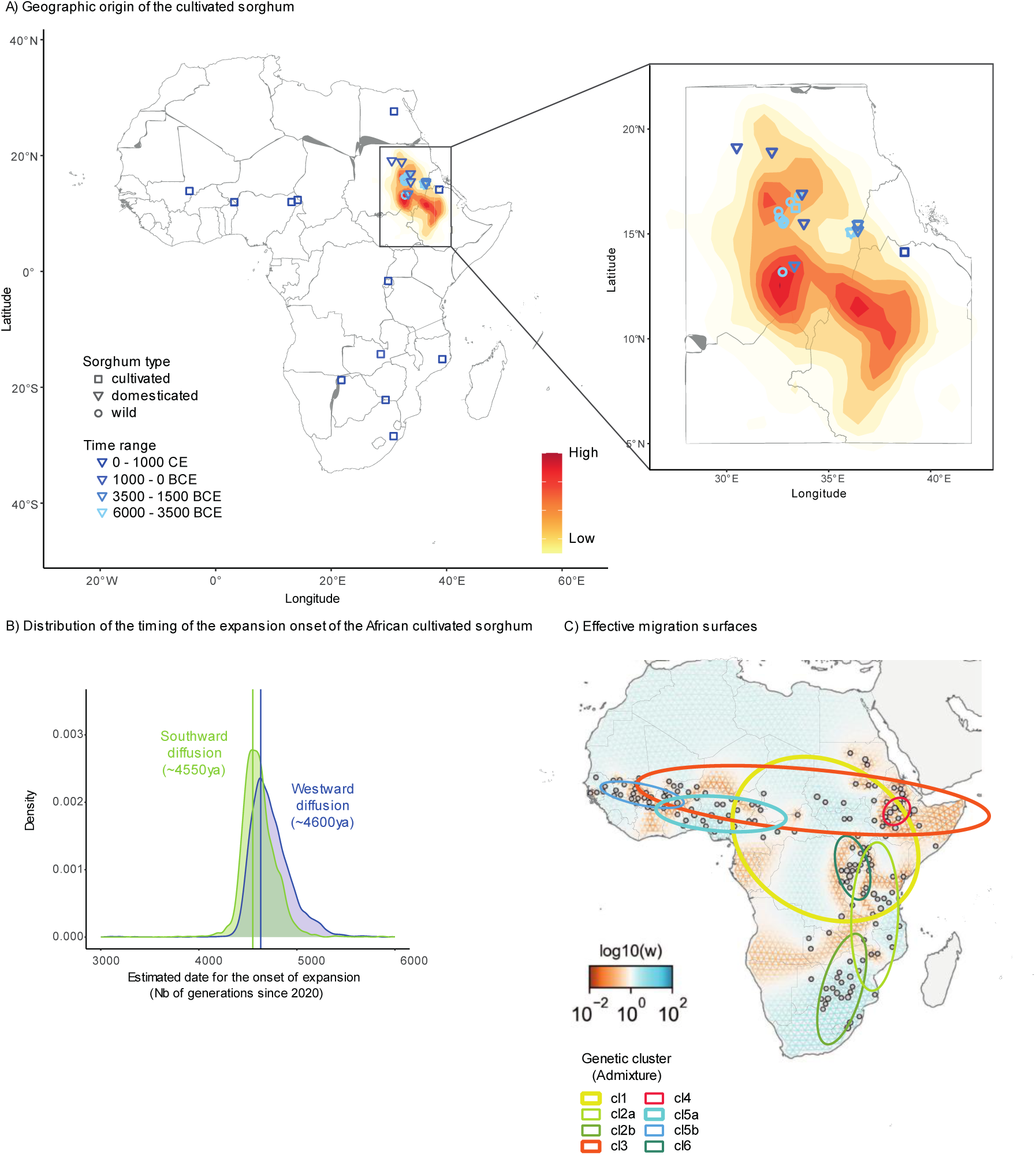
Geographic origin and diffusion of the African cultivated sorghum. A) Inferred geographic origin of the cultivated sorghum in Africa. The posterior distribution of the latitude and longitude of the origin in the spatial model was represented, the darker, the higher. The blue symbols correspond to the localizations of the archaeological sites in the inferred area of origin where the oldest remains of sorghum were found (from Table S4-B). B) Distribution of the timing of the beginning of the westward (blue) and southward (green) movements of diffusion of the African cultivated sorghum, removing the Ethiopian and the Egyptian archaeological sites from the analyses. C) Effective migration surfaces representing dispersal barriers (orange) and corridors (blue) without the cultivated accessions from the cluster cl5c. The geographic distribution of the eight genetic clusters considered in the analysis are illustrated on the FEEMS results map using standard deviational ellipses computed using the CrimeStat method implemented in the Standard Deviational Ellipse plugin for QGIS.

Using archaeobotanical remains to calibrate our model of spatial expansion (*Supplementary Information*, Table S2-B) led to an estimation for the onset of both the westward and southward diffusions of the African sorghum to have occurred around 4,600 ya (4,550 ya, 95% CI 4,340-4,979 for the southward diffusion and 4,600 ya, 95% CI 4,435-5,238 for the westward one; Fig. 2-B; *Supplementary Information*, SI Text section 2.2; Fig. S11).

The speed of the fronts of diffusion was estimated to be similar for the westward and southward diffusions, with a spread rate of 1.26 ± 0.4 km/year (80% CL: 0.67-1.85 km/year and r = 0.82.) and 1.14 ± 0.25 km/year (80% CL: 0.77-1.5 km/year, r = 0.88), respectively (*Supplementary Information*, Fig. S12-A).

Effective migration surfaces analysis using FEEMS (Marcus et al., 2021) mirrored the spatial organization of sorghum genetic diversity at different scales. Low migration rates were observed in Eastern Sudan and Western Ethiopia, the area of origin of the cultivated sorghum (Fig. 2-C). In this area, genetic differentiation between two cultivated accessions was higher than expected given their geographic distance. This suggests the existence of isolated populations (mainly clusters cl3 and cl4) between which gene flow was limited, while high diversity was observed within those two clusters. High genetic differentiation combined with high intra population diversity could result from low effective population sizes, and geographic and/or linguistic barriers to diffusion. Effective migration surfaces revealed other isolated populations in the African Great Lakes area of nilotic speakers (cluster cl6), in the South-Eastern Africa area of Guthrie S group speakers (cluster cl2b), and in the Nigeria area, where no Bantoid Benue-Congo speakers are located (mainly cluster cl5a).

## Discussion

In this study, we combined genomic, archaeobotanical and ethnolinguistic data to investigate the origin and spread of the cultivated sorghum in Africa. The origin of cultivated sorghum and the beginning of its expansion, as estimated by genomic data, converge with the result deduced from archaeological remains. The multidisciplinary approach adopted here indeed confirmed the East African origin of cultivated sorghum. The southward and westward expansions of sorghum were estimated to have begun at the same period, with a similar spread rate, by 4,600 BP, which is close to the period estimated from archaeological remains. The complex sorghum population genetic structure shows how its evolution dynamics were linked to both environmental and ethnolinguistic factors: effective migration surfaces analysis, mirroring the spatial organization of sorghum genetic diversity at different scales, confirmed that sorghum diffusion was constrained by geographic and/or ethnolinguistic barriers. Low migration rates, and high level of genetic diversity between and within genetic clusters in the area of origin, shed new light on the early period of sorghum cultivation.

Considering close associations between genetic clusters with present day ethnolinguistic groups in the area of origin and beyond, on one hand, and social barriers as anthropological characteristics on the other hand, allows upscaling our analysis over time and space.

### Geographic origin and expansion onset of African cultivated sorghum

Our results localized the onset of the expansion of the African cultivated sorghum in Eastern Sudan, specifically in the Southern Atbai region (the Eastern desert of Sudan), where the oldest domesticated sorghum remains are known (Fuller and Stevens, 2018). Besides the probabilistic map, one of the two hotspots of cultivated genetic diversity was localized in that same area (*Supplementary Information*, Fig. S-8), in line with the expectation of a higher diversity in the primary domestication center.

The second hotspot was observed in Western Africa and was mainly due to the accessions from the Western African cluster cl5c (guinea margaritiferum). The margaritiferum subrace is highly specific, both genetically (Burgarella et al., 2021) and morphologically, showing close relationships to wild sorghum with evidence of gene flow (Deu et al., 2006, 1995; Gilabert et al., 2023; Mace et al., 2013), and has been suggested to have originated from a secondary independent domestication event (Morris et al., 2013; Sagnard et al., 2011). This western hotspot could thus result from an independent sorghum domestication, cluster cl5c emerging from local Western African wild sorghum and/or from introgression between the wild and cultivated sorghum refueling the cultivated diversity with endemic wild one. More extended genomic study of this guinea margaritiferum group will certainly shed more light on its precise origin.

Calibrating with archaeobotanical data our genomic based spatial expansion model, we estimated that the westward and southward diffusions of cultivated sorghum from its center of origin in Eastern Africa occurred approximately 4,600 years ago (∼2,600 BCE). Our estimations are concomitant to the estimation for the cultivated pearl millet diffusion date (Burgarella et al., 2018), and correspond to a more ancient diffusion than the diffusion of the African rice and the African yam (Cubry et al., 2018; Scarcelli et al., 2019).The expansion of sorghum and pearl millet coincides with the intensification of the aridification of the “green Sahara”, and more specifically with the most severe drought episode the Sahara desert has experienced during the Holocene (Van der Meeren et al., 2022). The harsher conditions and reduced availability of Poaceae in the environment (Kröpelin et al., 2008) may have encouraged the cultivation of drought-resistant cereals such as sorghum and pearl millet, and the migration of people towards regions with more favorable conditions (Fuller et al., 2019).

It is worth noting that, with our approach, we estimated the onset of the expansion of cultivated sorghum, rather than the timing of the domestication process itself, which may largely predate the expansion. Even our estimation of the onset of expansion reflects a lower bound, given that the calibration is made on archaeobotanical remains that could be found only in the case of a rather important presence of the taxon, suggesting that reports of the presence of the crop post-date its actual arrival. The process of crop domestication is now understood as being protracted with the existence of an initial period of weak cultivation and/or management, rather than a rapid process accompanied by a domestication bottleneck (Allaby, 2010; Allaby et al., 2022, 2019; Fuller et al., 2014). The ABC results of our study do not support the hypothesis of a sharp population decline in the population of cultivated sorghum. Our finding is consistent with a study of modern and ancient sorghum genomes from Egyptian Nubia, which suggested that the loss of diversity observed in cultivated sorghum occurred gradually over time and was due to an accumulation of the mutation load rather than to a domestication bottleneck (Smith et al., 2019).

In line with these findings, sorghum archaeobotanical data argue in favor of the existence of a protracted period for sorghum in Africa. Charred remains of wild sorghum from huts and storage pits were identified in Egypt sites dating from approx. 8,000 BCE, and from potteries found in sites in Central Sudan active during the fifth and fourth millennium, suggesting that wild sorghum were likely collected and used in the Mid-Holocene (Fuller and Stevens, 2018; Lucarini and Radini, 2020; Wasylikowa and Dahlberg, 1999). The first evidence of cultivated sorghum was found in archaeological sites from Northern Sudan dating back to the 4^th^ millennium (Winchell et al., 2017) indicating that the domestication of sorghum was already ongoing at that time.

### Drivers of the genetic diversity of the African cultivated sorghum

The genetic diversity and structure of African sorghum have been suggested to be shaped by a combination of environmental, morphological (Billot et al., 2013; Deu et al., 2006; Lasky et al., 2015; Morris et al., 2013), and ethnolinguistic factors (Labeyrie et al., 2014; Westengen et al., 2014).

Morris et al. (2013) suggested that the diffusion of the cultivated sorghum has been driven by both geographic isolation and agroclimatic constraints. Similarly, we observed that some genetic clusters appeared to be preferentially distributed along a specific agroclimatic zone or altitudinal range, substantiating the role of climatic and environmental conditions in shaping sorghum diversity.

Analyzing a distinct minicore collection of African cultivated sorghum with simple sequence repeat markers, Westengen et al. (2014) identified three major sorghum groups that were co-distributed with the three main language families in Africa, in agreement with the farming/language dispersal hypothesis (Diamond and Bellwood, 2003). Our study, based on SNP markers, more finely divided sorghum diversity in nine groups and confirmed this previous finding of structuration of sorghum diversity at the language family scale, suggesting that sorghum genetic diversity is related to ethnolinguistic factors. An advantage of a fine description of sorghum genetic diversity is that we can associate sorghum diversity not only to a wide linguistic family level (Westengen et al., 2014) but also to a finer level and local linguistic groups, freeing us from potential correlations that could exist between language families and environmental variables at the higher level. This allows taking into consideration the ethnolinguistic groups that might have contributed to sorghum diversity diffusion or structure through the selection of sorghum locally-adapted to environmental conditions or usage.

The key role of social factors in structuring crop genetic diversity has been suggested to result from seed exchanges that were historically more oriented within than between sorghum populations (Westengen et al., 2014). Globally, this hypothesis, which was made explicit by Leclerc & Coppens d’Eeckenbrugge (2012), proposes that seed circulation is not random, and that speaking the same or a common language makes such interactions more likely. Linguistic organization of crop genetics diversity was observed for many crops, such maize (Orozco-Ramírez et al., 2016; Perales et al., 2005), pearl millet (Naino Jika et al., 2017) or fonio (Diop et al., 2023). In the case of fonio, farmers’ migrations are proposed as the mechanism leading to the fonio spatial structure in Senegal, which fits language diversity (Diop et al., 2023).

At a finer scale, local populations were detected in the area of origin of sorghum. Interestingly, those populations (clusters cl3 and cl4) are differentiated whereas they harbored a high level of intra-population diversity. This suggests that those populations were characterized by a low effective population size, and by limited gene flow between them, probably because of the existence of social barriers or adaptation to the environment or elevation. Action of social barriers on sorghum genetic diversity was well documented at the local scales (Labeyrie et al., 2016; Westengen et al., 2014). Notably, ethnolinguistic differentiation was also detected in the area of origin of cultivated sorghum in Northern Ethiopia in three sites with similar agro-ecological and topographic profiles, yet with admixture between the studied populations (Wendmu et al., 2023).

We identified three genetic clusters in the area of distribution of the Bantoid languages. The cluster cl6 was associated to the Guthrie zone J in the Great Lakes region, cluster cl2b to the zone S in South-Eastern Africa. These associations suggest that those sorghum populations moved with people, and since stayed associated with them. The cluster cl2a was associated with four Guthrie zones (F, G, N, P), which are located from north to south along a South-Eastern African corridor that reaches the area of distribution of the cluster cl2b, the cluster presenting the lowest level of genetic diversity. This could suggest a North to South migration route of sorghum (Fuller and Stevens, 2018). This is consistent with the two areas in Eastern and South-Eastern Africa with high effective migration rates. Using a similar analysis of effective migration rates of Bantu-speaking populations, congruent southeastern corridors along the coast from Tanzania to South Africa and the area of the Great Lakes were also detected for those human populations (Fortes-Lima et al., 2023).

Our estimations of the two fronts of dispersal of the cultivated African sorghum, at a rate of approx. 1 km/year, are consistent with previous estimations of the spread rate of agriculture in multiple areas of the world (reviewed in Fort (2021)). One striking result regarding the timing and the spread of the two movements of diffusion is that they appear to have occurred simultaneously. Dispersals along an east-west direction are expected to have happened more rapidly than those along a north-south axis (Diamond and Bellwood, 2003). Indeed, climates and environments show a higher variability latitudinally than longitudinally due to the fact that daylength and seasonality depends on the latitude only. Interestingly, a relatively constant movement rate through time was also suggested for Bantu-speaking populations, despite the variety of environments and interactions they had to have encountered during their migration (Fortes-Lima et al., 2023). The results obtained from Treemix suggest gene flow between cluster cl3, which is split between Eastern and Western Africa, and cluster cl2b in South-Eastern Africa. The interconnection between Eastern accessions, which are located in the sorghum area of origin, and cluster cl2b in South-Eastern Africa, suggests a route of diffusion that corresponds well to the one proposed by Fuller and Stevens (2018). The gene flow between cluster cl6 (Great Lakes region) and the widely distributed cluster cl1 could suggest a key role played by Bantu speakers of Guthrie zone J in sorghum diffusion from the area of origin and the Great Lakes region. The wide distribution of the cluster cl1 could suggest an independent diffusion, with secondary contact leading to signature of gene flow. The complexity of sorghum structuration could reflect the fact that African populations underwent multiple waves of migration during their history, known as the spread-over-spread model (Fortes-Lima et al., 2023; Seidensticker et al., 2021).

## Conclusion

Combining genomics and archaeobotanical data, this study corroborates the Eastern African origin of the cultivated sorghum. It also proposes two simultaneous and synchronous fronts of diffusions from the area of origin towards the West and the South. Overall, our results also confirm the complex geographic-based genetic structuring of the cultivated sorghum, whose genetic structure at the African continental scale appears to be shaped by a combination of multiple factors, environmental and ethnolinguistic ones, acting at different temporal and spatial scales. One major outstanding question on the evolutionary history of cultivated plants and the history of their domestication relies on their origin and how they dispersed from there: have crops dispersed through a demic or cultural diffusion, or a combination of the two (Hewlett et al., 2002; Leclerc and Coppens d’Eeckenbrugge, 2012Under a scenario of a demic diffusion, one would expect crops to be structured by social factors as they move together with the farmers while a cultural diffusion model would lead to patterns of isolation by distance since it depends on interactions with neighboring societies. If our study helps the understanding of the beginning of sorghum domestication, a comparative study of population genetics of ancient human and crop populations (see for example Tao et al. (2023)) and/or additional archaeobotanical data (see for example Cobo et al. (2019)) would be necessary to give insight into the main mechanism, a demic and/or a cultural diffusion, responsible for the diffusion of agriculture, and thus help us to better understand their evolutionary history.

## Data and code availability

The raw GBS data were available in the NCBI Sequence Read Archive from references Brenton et al., 2016; Lasky et al., 2015; Yu et al., 2016. The vcf file containing the filtered SNPs and the passport data of the accessions analyzed in the study are publicly available on the CIRAD Dataverse repository, under the doi identifier https://doi.org/10.18167/DVN1/SAAMUT. The R scripts for the analyses performed in the study are available on the CIRAD gitlab plateform (https://gitlab.cirad.fr/agap/sorgho/africrop_sorghum).

## Supplemental material

Supplementary material is available online at: https://www.biorxiv.org/content/10.1101/2025.04.16.648676v1.supplementary-material

## Acknowledgments

This work was supported by an ANR grant (AfriCrop project, ANR-13-BSV7-0017). We acknowledge the following bioinformatic facilities for providing HPC resources and support: the Core Cluster of the Institut Français de Bioinformatique, the MESO@LR-Platform at the University of Montpellier and the INRAE MIGALE bioinformatics facility (MIGALE, INRAE, 2020. Migale bioinformatics Facility, doi: 10.15454/1.5572390655343293E12). The preprint version 1 has been peer-reviewed and recommended by Peer Community in Evolutionary Biology (https://doi.org/10.24072/pci.evolbiol.100865).

## References

1. Alexander, D.H., Novembre, J., Lange, K., 2009. Fast model-based estimation of ancestry in unrelated individuals. Genome Res. 19, 1655–1664. 10.1101/gr.094052.109

2. Allaby, R., 2010. Integrating the processes in the evolutionary system of domestication. J. Exp. Bot. 61, 935–944. 10.1093/jxb/erp382

3. Allaby, R.G., Stevens, C.J., Kistler, L., Fuller, D.Q., 2022. Emerging evidence of plant domestication as a landscape-level process. Trends Ecol. Evol. 37, 268–279. 10.1016/j.tree.2021.11.002

4. Allaby, R.G., Ware, R.L., Kistler, L., 2019. A re-evaluation of the domestication bottleneck from archaeogenomic evidence. Evol. Appl. 12, 29–37. 10.1111/eva.12680

5. Beck, H.E., Zimmermann, N.E., McVicar, T.R., Vergopolan, N., Berg, A., Wood, E.F., 2018. Present and future Köppen-Geiger climate classification maps at 1-km resolution. Sci. Data 5, 180214. 10.1038/sdata.2018.214

6. Billot, C., Ramu, P., Bouchet, S., Chantereau, J., Deu, M., Gardes, L., Noyer, J.-L., Rami, J.-F., Rivallan, R., Li, Y., Lu, P., Wang, T., Folkertsma, R.T., Arnaud, E., Upadhyaya, H.D., Glaszmann, J.-C., Hash, C.T., 2013. Massive sorghum collection genotyped with SSR markers to enhance use of global genetic resources. PLOS ONE 8, e59714. 10.1371/journal.pone.0059714

7. Bolger, A.M., Lohse, M., Usadel, B., 2014. Trimmomatic: a flexible trimmer for Illumina sequence data. Bioinformatics 30, 2114–2120. 10.1093/bioinformatics/btu170

8. Brenton, Z.W., Cooper, E.A., Myers, M.T., Boyles, R.E., Shakoor, N., Zielinski, K.J., Rauh, B.L., Bridges, W.C., Morris, G.P., Kresovich, S., 2016. A genomic resource for the development, improvement, and exploitation of sorghum for bioenergy. Genetics 204, 21–33. 10.1534/genetics.115.183947

9. Burgarella, C., Berger, A., Glémin, S., David, J., Terrier, N., Deu, M., Pot, D., 2021. The road to sorghum domestication: evidence from nucleotide diversity and gene expression patterns. Front. Plant Sci. 12. 10.3389/fpls.2021.666075.

10. Burgarella, C., Cubry, P., Kane, N.A., Varshney, R.K., Mariac, C., Liu, X., Shi, C., Thudi, M., Couderc, M., Xu, X., Chitikineni, A., Scarcelli, N., Barnaud, A., Rhoné, B., Dupuy, C., François, O., Berthouly-Salazar, C., Vigouroux, Y., 2018. A western Sahara centre of domestication inferred from pearl millet genomes. Nat. Ecol. Evol. 2, 1377–1380. 10.1038/s41559-018-0643-y

11. Chang, C.C., Chow, C.C., Tellier, L.C., Vattikuti, S., Purcell, S.M., Lee, J.J., 2015. Second-generation PLINK: rising to the challenge of larger and richer datasets. GigaScience 4, s13742-015-0047–8. 10.1186/s13742-015-0047-8

12. Cobo, J.M., Fort, J., Isern, N., 2019. The spread of domesticated rice in eastern and southeastern Asia was mainly demic. J. Archaeol. Sci. 101, 123–130. 10.1016/j.jas.2018.12.001

13. Csilléry, K., François, O., Blum, M.G.B., 2012. abc: an R package for approximate Bayesian computation (ABC). Methods Ecol. Evol. 3, 475–479. 10.1111/j.2041-210X.2011.00179.x

14. Cubry, P., Tranchant-Dubreuil, C., Thuillet, A.-C., Monat, C., Ndjiondjop, M.-N., Labadie, K., Cruaud, C., Engelen, S., Scarcelli, N., Rhoné, B., Burgarella, C., Dupuy, C., Larmande, P., Wincker, P., François, O., Sabot, F., Vigouroux, Y., 2018. The rise and fall of African rice cultivation revealed by analysis of 246 new genomes. Curr. Biol. 28, 2274–2282.e6. 10.1016/j.cub.2018.05.066

15. Currat, M., Ray, N., Excoffier, L., 2004. splatche: a program to simulate genetic diversity taking into account environmental heterogeneity. Mol. Ecol. Notes 4, 139–142. 10.1046/j.1471-8286.2003.00582.x

16. Danecek, P., Auton, A., Abecasis, G., Albers, C.A., Banks, E., DePristo, M.A., Handsaker, R.E., Lunter, G., Marth, G.T., Sherry, S.T., McVean, G., Durbin, R., 1000 Genomes Project Analysis Group, 2011. The variant call format and VCFtools. Bioinformatics 27, 2156– 2158. 10.1093/bioinformatics/btr330

17. de Wet, J.M.J., Huckabay, J.P., 1967. The origin of *Sorghum bicolor*. II. Distribution and domestication. Evolution 21, 787–802. 10.2307/2406774

18. DePristo, M.A., Banks, E., Poplin, R., Garimella, K.V., Maguire, J.R., Hartl, C., Philippakis, A.A., del Angel, G., Rivas, M.A., Hanna, M., McKenna, A., Fennell, T.J., Kernytsky, A.M., Sivachenko, A.Y., Cibulskis, K., Gabriel, S.B., Altshuler, D., Daly, M.J., 2011. A framework for variation discovery and genotyping using next-generation DNA sequencing data. Nat. Genet. 43, 491–498. 10.1038/ng.806

19. Deu, M., Hamon, P., Dufour, P., D’ ont, A., Lanaud, C., Chantereau, J., 1995. Mitochondrial DNA diversity in wild and cultivated sorghum. Genome 38, 635–645. 10.1139/g95-081

20. Deu, M., Rattunde, F., Chantereau, J., 2006. A global view of genetic diversity in cultivated sorghums using a core collection. Genome 49, 168–180. 10.1139/g05-092

21. Diamond, J., Bellwood, P., 2003. Farmers and their languages: the first expansions. Science 300, 597–603. 10.1126/science.1078208

22. Dinno A (2024). dunn.test: Dunn’s Test of Multiple Comparisons Using Rank Sums. R package version 1.3.6. Available at https://cran.r-project.org/web/packages/dunn.test/

23. Diop, B.M., Guèye, M.C., Leclerc, C., Deu, M., Zekraoui, L., Calatayud, C., Rivallan, R., Kaly, J.R., Cissé, M., Piquet, M., Diack, O., Ngom, A., Berger, A., Ndoye, I., Ndir, K., Vigouroux, Y., Kane, N.A., Barnaud, A., Billot, C., 2023. Ethnolinguistic and genetic diversity of fonio (*Digitaria exilis*) in Senegal. Plants People Planet. 10.1002/ppp3.10428

24. Evanno, G., Regnaut, S., Goudet, J., 2005. Detecting the number of clusters of individuals using the software structure: a simulation study. Mol. Ecol. 14, 2611–2620. 10.1111/j.1365-294X.2005.02553.x

25. FAO, 2023. Agricultural production statistics 2000–2022. https://openknowledge.fao.org/handle/20.500.14283/cc9205en

26. Fitak, R.R., 2021. OptM: estimating the optimal number of migration edges on population trees using Treemix. Biol. Methods Protoc. 6, bpab017. 10.1093/biomethods/bpab017

27. Fort, J., 2021. The spread of agriculture: quantitative laws in Prehistory?, in: Pardo-Gordó, S., Bergin, S. (Eds.), Simulating Transitions to Agriculture in Prehistory, Computational Social Sciences. Springer International Publishing, Cham, pp. 17–28. 10.1007/978-3-030-83643-6_2

28. Fort, J., Pujol, T., Cavalli-Sforza, L.L., 2004. Palaeolithic populations and waves of advance. Camb. Archaeol. J. 14, 53–61. 10.1017/S0959774304000046

29. Fortes-Lima, C.A., Burgarella, C., Hammarén, R., Eriksson, A., Vicente, M., Jolly, C., Semo, A., Gunnink, H., Pacchiarotti, S., Mundeke, L., Matonda, I., Muluwa, J.K., Coutros, P., Nyambe, T.S., Cikomola, J.C., Coetzee, V., de Castro, M., Ebbesen, P., Delanghe, J., Stoneking, M., Barham, L., Lombard, M., Meyer, A., Steyn, M., Malmström, H., Rocha, J., Soodyall, H., Pakendorf, B., Bostoen, K., Schlebusch, C.M., 2023. The genetic legacy of the expansion of Bantu-speaking peoples in Africa. Nature 1–8. 10.1038/s41586-023-06770-6

30. Fuller, D., Champion, L., Stevens, C., 2019. Comparing the tempo of cereal dispersal and the agricultural transition: two African and one West Asian trajectory, in: Eichhorn, B., Hohn, A. (Eds.), In: Eichhorn, B and Hohn, A, (Eds.) Trees, Grasses and Crops – People and Plants in Sub-Saharan Africa and Beyond. Verlag Dr. Rudolf Habelt GmbH: Bonn, Germany. (2019). Verlag Dr. Rudolf Habelt GmbH, Bonn, Germany.

31. Fuller, D.Q., Denham, T., Arroyo-Kalin, M., Lucas, L., Stevens, C.J., Qin, L., Allaby, R.G., Purugganan, M.D., 2014. Convergent evolution and parallelism in plant domestication revealed by an expanding archaeological record. Proc. Natl. Acad. Sci. 111, 6147–6152. 10.1073/pnas.1308937110

32. Fuller, D.Q., Stevens, C.J., 2018. Sorghum domestication and diversification: a current archaeobotanical perspective, in: Mercuri, A.M., D’Andrea, A.C., Fornaciari, R., Höhn, A. (Eds.), Plants and People in the African Past: Progress in African Archaeobotany. Springer International Publishing, Cham, pp. 427–452. 10.1007/978-3-319-89839-1_19

33. Gilabert, A., Burgarella, C., Calatayud, C., Berger, A., Rami, J.-F., Pot, D., Deu, M., 2023. Shedding light on the evolutionary history of wild and cultivated african sorghum: the guinea margaritiferum case. Poster presented at: Sorghum in the 21st century: Resiliency and sustainability in the face of climate change; June 5-9 2023; Montpellier, France.

34. Goudet, J., 2005. hierfstat, a package for R to compute and test hierarchical *F*-statistics. Mol. Ecol. Notes 5, 184–186. 10.1111/j.1471-8286.2004.00828.x

35. Guthrie, M., 1971. Comparative Bantu: An Introduction to the Comparative Linguistics and Prehistory of the Bantu Languages. Gregg, Famborough, UK.

36. Guthrie, M., 1948. The Classification of the Bantu languages. Oxford University Press.

37. Harlan, J.R., Stemler, A.B.L., 1976. The races of sorghum in Africa, in: Harlan, J.R., Wet, J.M.J. de, Stemler, A.B.L. (Eds.), Origins of African Plant Domestication. De Gruyter Mouton, Berlin, New York, pp. 465–478. doi:10.1515/9783110806373.465

38. Hewlett, B.S., De Silvestri, A., Guglielmino, C.R., 2002. Semes and genes in Africa. Curr. Anthropol. 43, 313–321. 10.1086/339379

39. Jerardino, A., Fort, J., Isern, N., Rondelli, B., 2014. Cultural diffusion was the main driving mechanism of the Neolithic transition in Southern Africa. PLOS ONE 9, e113672. 10.1371/journal.pone.0113672

40. Kimber, C.T., 2000. Origins of domesticated sorghum and its early diffusion to India and China, in: Sorghum: Origin, History, Technology, and Production. John Wiley & Sons, New York, pp. 3–98.

41. Kröpelin, S., Verschuren, D., Lézine, A.-M., Eggermont, H., Cocquyt, C., Francus, P., Cazet, J.-P., Fagot, M., Rumes, B., Russell, J.M., Darius, F., Conley, D.J., Schuster, M., von Suchodoletz, H., Engstrom, D.R., 2008. Climate-driven ecosystem succession in the Sahara: The past 6000 Years. Science 320, 765–768. 10.1126/science.1154913

42. Labeyrie, V., Deu, M., Barnaud, A., Calatayud, C., Buiron, M., Wambugu, P., Manel, S., Glaszmann, J.-C., Leclerc, C., 2014. Influence of ethnolinguistic diversity on the sorghum genetic patterns in subsistence farming systems in Eastern Kenya. PLOS ONE 9, e92178. 10.1371/journal.pone.0092178

43. Labeyrie, V., Thomas, M., Muthamia, Z.K., Leclerc, C., 2016. Seed exchange networks, ethnicity, and sorghum diversity. Proc. Natl. Acad. Sci. 113, 98–103. 10.1073/pnas.1513238112

44. Lasky, J.R., Upadhyaya, H.D., Ramu, P., Deshpande, S., Hash, C.T., Bonnette, J., Juenger, T.E., Hyma, K., Acharya, C., Mitchell, S.E., Buckler, E.S., Brenton, Z., Kresovich, S., Morris, G.P., 2015. Genome-environment associations in sorghum landraces predict adaptive traits. Sci. Adv. 1, e1400218. 10.1126/sciadv.1400218

45. Leclerc, C., Coppens d’Eeckenbrugge, G., 2012. Social organization of crop genetic diversity. The G × E × S interaction model. Diversity 4, 1–32. 10.3390/d4010001

46. Leinonen, R., Sugawara, H., Shumway, M., on behalf of the International Nucleotide Sequence Database Collaboration, 2011. The Sequence Read Archive. Nucleic Acids Res. 39, D19–D21. 10.1093/nar/gkq1019

47. Lemaire, L., Jay, F., Lee, I.-H., Csilléry, K., Blum, M.G.B., 2016. Goodness-of-fit statistics for approximate Bayesian computation. 10.48550/arXiv.1601.04096

48. Lewis, M., 2009. Ethnologue: languages of the World, 16th edition. SIL International.

49. Li, H., Durbin, R., 2009. Fast and accurate short read alignment with Burrows–Wheeler transform. Bioinformatics 25, 1754–1760. 10.1093/bioinformatics/btp324

50. Li, H., Handsaker, B., Wysoker, A., Fennell, T., Ruan, J., Homer, N., Marth, G., Abecasis, G., Durbin, R., 1000 Genome Project Data Processing Subgroup, 2009. The Sequence Alignment/Map format and SAMtools. Bioinformatics 25, 2078–2079. 10.1093/bioinformatics/btp352

51. Lucarini, G., Radini, A., 2020. First direct evidence of wild plant grinding process from the Holocene Sahara: use-wear and plant micro-residue analysis on ground stone tools from the Farafra asis, Egypt. Quat. Int., Africa under the microscope: What’s new in techno-functional analyses in archaeology? 555, 66–84. 10.1016/j.quaint.2019.07.028

52. Mace, E.S., Tai, S., Gilding, E.K., Li, Y., Prentis, P.J., Bian, L., Campbell, B.C., Hu, W., Innes, D.J., Han, X., Cruickshank, A., Dai, C., Frère, C., Zhang, H., Hunt, C.H., Wang, X., Shatte, T., Wang, M., Su, Z., Li, J., Lin, X., Godwin, I.D., Jordan, D.R., Wang, J., 2013. Whole-genome sequencing reveals untapped genetic potential in Africa’s indigenous cereal crop sorghum. Nat. Commun. 4, 2320. 10.1038/ncomms3320

53. Maho, J.F., 2009. NUG nline : the online version of the ew Updated Guthrie ist, a referential classification of the Bantu languages.

54. Marcus, J., Ha, W., Barber, R.F., Novembre, J., 2021. Fast and flexible estimation of effective migration surfaces. eLife 10, e61927. 10.7554/eLife.61927

55. McCormick, R.F., Truong, S.K., Sreedasyam, A., Jenkins, J., Shu, S., Sims, D., Kennedy, M., Amirebrahimi, M., Weers, B.D., McKinley, B., Mattison, A., Morishige, D.T., Grimwood, J., Schmutz, J., Mullet, J.E., 2018. The *Sorghum bicolor* reference genome: improved assembly, gene annotations, a transcriptome atlas, and signatures of genome organization. Plant J. 93, 338–354. 10.1111/tpj.13781

56. Morris, G.P., Ramu, P., Deshpande, S.P., Hash, C.T., Shah, T., Upadhyaya, H.D., Riera-Lizarazu, O., Brown, P.J., Acharya, C.B., Mitchell, S.E., Harriman, J., Glaubitz, J.C., Buckler, E.S., Kresovich, S., 2013. Population genomic and genome-wide association studies of agroclimatic traits in sorghum. Proc. Natl. Acad. Sci. 110, 453–458. 10.1073/pnas.1215985110

57. Naino Jika, A.K., Dussert, Y., Raimond, C., Garine, E., Luxereau, A., Takvorian, N., Djermakoye, R.S., Adam, T., Robert, T., 2017. Unexpected pattern of pearl millet genetic diversity among ethno-linguistic groups in the Lake Chad Basin. Heredity 118, 491–502. 10.1038/hdy.2016.128

58. Orozco-Ramírez, Q., Ross-Ibarra, J., Santacruz-Varela, A., Brush, S., 2016. Maize diversity associated with social origin and environmental variation in Southern Mexico. Heredity 116, 477–484. 10.1038/hdy.2016.10

59. Perales, H.R., Benz, B.F., Brush, S.B., 2005. Maize diversity and ethnolinguistic diversity in Chiapas, Mexico. Proc. Natl. Acad. Sci. 102, 949–954. 10.1073/pnas.0408701102

60. Perrier, X., Jacquemoud-Collet, J.-P., 2019. DARwin - Dissimilarity Analysis and Representation for Windows. Available at: https://darwin.cirad.fr/

61. Pickrell, J.K., Pritchard, J.K., 2012. Inference of population splits and mixtures from genome-wide allele frequency data. PLOS Genet. 8, e1002967. 10.1371/journal.pgen.1002967

62. Pritchard, J.K., Stephens, M., Donnelly, P., 2000. Inference of population structure using multilocus genotype data. Genetics 155, 945–959. 10.1093/genetics/155.2.945

63. Purcell, S., Neale, B., Todd-Brown, K., Thomas, L., Ferreira, M.A.R., Bender, D., Maller, J., Sklar, P., Bakker, P.I.W. de, Daly, M.J., Sham, P.C., 2007. PLINK: a tool set for whole-genome association and population-based linkage analyses. Am. J. Hum. Genet. 81, 559–575. 10.1086/519795

64. R Core Team, 2024. R: A Language and Environment for Statistical Computing. R Foundation for Statistical Computing, Vienna, Austria. URL https://www.R-project.org/

65. Sagnard, F., Deu, M., Dembélé, D., Leblois, R., Touré, L., Diakité, M., Calatayud, C., Vaksmann, M., Bouchet, S., Mallé, Y., Togola, S., Traoré, P.C.S., 2011. Genetic diversity, structure, gene flow and evolutionary relationships within the *Sorghum bicolor* wild–weedy–crop complex in a western African region. Theor. Appl. Genet. 123, 1231–1246. 10.1007/s00122-011-1662-0

66. Saitou, N., Nei, M., 1987. The neighbor-joining method: a new method for reconstructing phylogenetic trees. Mol. Biol. Evol. 4, 406–425. 10.1093/oxfordjournals.molbev.a040454

67. Scarcelli, N., Cubry, P., Akakpo, R., Thuillet, A.-C., Obidiegwu, J., Baco, M.N., Otoo, E., Sonké, B., Dansi, A., Djedatin, G., Mariac, C., Couderc, M., Causse, S., Alix, K., Chaïr, H., François, O., Vigouroux, Y., 2019. Yam genomics supports West Africa as a major cradle of crop domestication. Sci. Adv. 5, eaaw1947. 10.1126/sciadv.aaw1947

68. Seidensticker, D., Hubau, W., Verschuren, D., Fortes-Lima, C., de Maret, P., Schlebusch, C.M., Bostoen, K., 2021. Population collapse in Congo rainforest from 400 CE urges reassessment of the Bantu expansion. Sci. Adv. 7, eabd8352. 10.1126/sciadv.abd8352

69. Smith, C.W., Frederiksen, R.A., 2000. Sorghum: origin, history, technology, and production, Wiley Series in Crop Science. Wiley.

70. Smith, O., Nicholson, W.V., Kistler, L., Mace, E., Clapham, A., Rose, P., Stevens, C., Ware, R., Samavedam, S., Barker, G., Jordan, D., Fuller, D.Q., Allaby, R.G., 2019. A domestication history of dynamic adaptation and genomic deterioration in Sorghum. Nat. Plants 5, 369–379. 10.1038/s41477-019-0397-9

71. Tao, L., Yuan, H., Zhu, K., Liu, X., Guo, J., Min, R., He, H., Cao, D., Yang, X., Zhou, Z., Wang, R., Zhao, D., Ma, H., Chen, J., Zhao, J., Li, Y., He, Y., Suo, D., Zhang, R., Li, S., Li, L., Yang, F., Li, H., Zhang, L., Jin, L., Wang, C.-C., 2023. Ancient genomes reveal millet farming-related demic diffusion from the Yellow River into southwest China. Curr. Biol. 0. 10.1016/j.cub.2023.09.055

72. Van der Meeren, T., Verschuren, D., Sylvestre, F., Nassour, Y.A., Naudts, E.L., Aguilar Ortiz, L.E., Deschamps, P., Tachikawa, K., Bard, E., Schuster, M., Abderamane, M., 2022. A predominantly tropical influence on late Holocene hydroclimate variation in the hyperarid central Sahara. Sci. Adv. 8, eabk1261. 10.1126/sciadv.abk1261

73. Wasylikowa, K., Dahlberg, J., 1999. Sorghum in the economy of the early Neolithic nomadic tribes at Nabta Playa, Southern Egypt, in: van der Veen, M. (Ed.), The Exploitation of Plant Resources in Ancient Africa. Springer US, Boston, MA, pp. 11–31. 10.1007/978-1-4757-6730-8_2

74. Wendmu, T.A., Gebrelibanos, T.S., Kovi, M.R., Ring, K.H., de Boer, H.J., Abera, F.A., Westengen, O.T., 2023. “ People gathered by sorghum”: cultural practices and sorghum diversity in Northern Ethiopia. Hum. Ecol. 51, 923–935. 10.1007/s10745-023-00442-9

75. Westengen, O.T., Okongo, M.A., Onek, L., Berg, T., Upadhyaya, H., Birkeland, S., Kaur Khalsa, S.D., Ring, K.H., Stenseth, N.C., Brysting, A.K., 2014. Ethnolinguistic structuring of sorghum genetic diversity in Africa and the role of local seed systems. Proc. Natl. Acad. Sci. 111, 14100–14105. 10.1073/pnas.1401646111

76. Winchell, F., Stevens, C.J., Murphy, C., Champion, L., Fuller, D.Q., 2017. Evidence for sorghum domestication in fourth millennium BC Eastern Sudan: spikelet morphology from ceramic impressions of the Butana group. Curr. Anthropol. 58, 673–683. 10.1086/693898

77. Yu, X., Li, X., Guo, T., Zhu, C., Wu, Y., Mitchell, S.E., Roozeboom, K.L., Wang, D., Wang, M.L., Pederson, G.A., Tesso, T.T., Schnable, P.S., Bernardo, R., Yu, J., 2016. Genomic prediction contributing to a promising global strategy to turbocharge gene banks. Nat. Plants 2, 1–7. 10.1038/nplants.2016.150

