## Supplementary material for "Integrating ethnolinguistic and archaeobotanical data to uncover the origin and dispersal of cultivated sorghum in Africa: a genomic perspective"

### Table of Contents:

| | |
|---|---|
| Supplementary results | Page 2 |
| Figure S1: Demographic scenarios | Page 6 |
| Figure S2: Numbers of genetic clusters | Page 7 |
| Figure S3: Sorghum genetic clusters | Page 8 |
| Figure S4: Genetic clusters and botanical races | Page 9 |
| Figure S5: Principal Component Analysis | Page 10 |
| Figure S6: Neighbor-joining tree | Page 12 |
| Figure S7: Treemix results | Page 13 |
| Figure S8: Hotspots of genetic diversity | Page 14 |
| Figure S9: Genetic clusters, agroclimatic zones and ethnolinguistic diversity | Page 15 |
| Figure S10: Geographic origin of the African cultivated sorghum | Page 18 |
| Figure S11: Timing of the onset of African cultivated sorghum expansion | Page 23 |
| Figure S12: Speed of the fronts of diffusion | Page 24 |
| Table S1: Splatche parameters | Page 25 |
| Table S2: Archaeological sites | Page 27 |
| Table S3: Genetic diversity | Page 29 |
| Table S4: Genetic clusters, agroclimatic zones and ethnolinguistic diversity | Page 30 |
| Table S5: Goodness-of-fit of the ABC models | Page 32 |
| Supplementary references | Page 33 |

### Supplementary results

#### *1. Population genetic structure*

#### *1.1 Cluster analysis*

We first assessed the population structure of the African cultivated sorghums using ADMIXTURE (Alexander et al., 2009), which implements a model-based approach to estimate individuals' ancestries. We used the dataset filtered for high quality SNPs. We ran the program for a number of clusters *K* ranging from 1 to 20, with 10 replicates for each value of *K*. For each value of *K*, we computed the mean and standard deviation of the cross-validation (CV) errors over the 10 replicates. The plot of the cross-validation errors according to the number of genetic clusters clearly showed a minimal CV errors value for nine clusters indicating that the optimal number of clusters in our sampling was nine (*Supplementary Information*, Fig. S2). The supplementary figure S3 illustrates for each accession the probabilities of ancestry for each cluster, the number of clusters ranging from 2 to 10. For *K* = 2, the analysis differentiated a group of accessions almost exclusively found in Western Africa (cluster formed by the hereafter clusters cl5a, b and c). At *K*=3, the cultivated accessions were subdivided into three clusters, the Western African cluster cl5, one cluster that included accessions from Eastern and Western Africa (cl3 & 4) and the last one with a wide geographic repartition (cl1, 2a, 2b and 6). At *K*=4 and K=5, the westernmost accessions (cl5c) and the Southeastern accessions (cl2) separated from the cl5 group and the widest group, respectively. At *K*=5, the aforementioned cluster from Eastern and Western Africa exclusively included here the Eastern African accessions that will be assigned to cluster cl4. The remaining accessions were found to be admixed between cl4 and the wide cluster grouping clusters cl1 and cl6. Increasing the numbers of clusters from 6 to 8 led to the distinction of the clusters cl6, cl3 and the differentiation of the clusters cl5a and cl5b. Lastly, at *K*=9 the cluster cl2 split into two clusters, the cluster 2b being restricted to southeastern Africa while the 2a was distributed in Southern and Eastern Africa.

#### *1.2 Principal Component Analysis*

Population structure was also assessed using a Principal Component Analysis (PCA) to visualize the genetic proximity between the cultivated accessions of sorghum. The first three axes of the PCA explained 55.4% of the total variance. The first axis differentiated the Western African accessions that were assigned to the cluster cl5c with the ADMIXTURE analysis (*Supplementary Information*, Fig. S5-A and B). Noteworthy, using whole genome sequence data and a distinct sampling, we observed that the accessions of this cluster cl5c in common between the two analyses clustered together with accessions clearly identified as margaritiferum accessions following the taxonomic classification proposed by Snowden (1936; results not shown; Mendy et al., 2023). The margaritiferum subrace is highly specific, both genetically (Burgarella et al., 2021) and morphologically, showing close relationships to wild sorghum with evidence of gene flow (Deu et al., 2006, 1995; Gilabert et al., 2023; Mace et al., 2013), and has been suggested to have originated from a secondary independent domestication event (Morris et al., 2013; Sagnard et al., 2011). Along the second axis, two clusters could be distinguished from the other accessions: the clusters cl4 and cl3 (*Supplementary Information*, Fig. S5-A and C). The third axis allowed to differentiate the Western African from the ADMIXTURE clusters cl5a and 5b from the other accessions (*Supplementary Information*, Fig. S5-B and C).

#### *1.3 Associations with agroclimatic zones and ethnolinguistic groups*

To better understand the geographic distribution of the genetic clusters, we looked for associations between these groups and elevation or agroclimatic and ethnolinguistic zones. For the agroclimatic zones, we used the Köppen-Geiger climate classification, which defines climate zones with different types of vegetation according to the temperature and/or the dryness (Beck et al., 2018), and for the ethnolinguistic zones, the language classification referenced in the database Ethnologue Version 16 (Lewis, 2009). A total of 73% of our accessions were in two agroclimatic zones: the tropical savanna zone (Aw) characterized by a tropical climate with winter dry season, and the semi-arid climate zone (Bsh) characterized by a hot steppe climate (*Supplementary Information*, Fig. S9-A and B, Table S4-A). Most of the genetic clusters were distributed in several agroclimatic zones but three of them appeared to be preferentially found in a specific zone: 79% and 67% of the accessions of the clusters cl5a (23 of 29 accessions) and cl6 (16 of 24) were distributed in a tropical savanna climate (Aw zone), and 60% of the accessions of the cluster cl2b were in a hot semi-arid climate (Bsh zone).

As for the linguistics, when considering the three main language families observed in Africa, the Afro-Asiatic, the Nilo-Saharan and the Niger-Congo families, the latter being subdivided into the Atlantic-Congo and the Mande, we observed that two genetic clusters (cl2a and cl2b) seemed preferentially found in areas from the Atlantic-Congo family, which contained more than 95% of their accessions, two clusters, cl3 and cl4, were co-distributed with the Afro-Asiatic family, with respectively 86 and 100% of their accessions being found in areas from the Afro-Asiatic group (*Supplementary Information*, Table S4-B; Fig. 1-B and S9-D). The analysis of the chi-squared residuals confirmed those associations and also indicate that the clusters cl1 and cl6 were associated with the Nilo-Saharan family. When considering a finer language classification of the Atlantic-Congo language family, we observed that apart from two Guthrie linguistic zones, the Great Lake zone J and the southeastern zone S, for which respectively 94% (16 of 17) and 78% (21 of 27) of the accessions belonged to one genetic cluster (clusters cl6 and cl2b respectively), none of the other linguistic zones was associated to a single genetic cluster when multiple accessions were present (Fig.1-C; *Supplementary Information*, Fig. S9-D and Table S4-B). Yet, four clusters showed a distribution associated with a linguistic zone: all of the 17 accessions of the cl4 and 12 of the 14 accessions of the cluster cl3 (86%) were distributed in areas of the Afro-Asiatic language family, 21 of the 22 accessions of the cluster cl2b (95%) were from the Guthrie linguistic zone S and 16 of the 24 accessions of the cluster cl6 (67%) were from the Guthrie linguistic zone J (Fig. 1-C;

*Supplementary Information*, Fig. S9-D and Table S4-B). The analysis of the chi-squared residuals revealed that cluster cl2b was indeed strongly (Chi2 residual over 9) associated with the Narrow Bantu Guthrie linguistic zone S and that the cluster cl6, still associated with Nilo-saharan languages, was strongly associated with the Narrow Bantu Guthrie linguistic zone J. In addition, the cluster cl2a was associated with four geographically close Bantu Guthrie linguistic zones (zones F, G, N and P). Finally, the westernmost clusters cl5b and cl5c were associated with the Atlantic-Congo and the Mande, the cluster cl5b being also associated with the Volta-Congo linguistic group. The cluster cl5a was associated with multiple Niger-Congo, non-Bantu, groups and with the Afro-Asiatic Chadic group.

### *2. Geographic origin and barriers to dispersal for the cultivated sorghum*

#### *2.1 Geographic origin of the cultivated sorghum*

We inferred the geographic origin of the cultivated African sorghum using spatially-explicit simulations (Currat et al., 2004). The previous population genetics analyses of our African sample highlighted the genetic cluster cl5c as a peculiar cluster, represented by accessions from the guinea margaritiferum group. This peculiar status of the maragaritiferum accessions may result from introgression with wild populations or from an independent domestication origin (see Materials and Methods and results), and led us to exclude them for these analyses. Yet, we provide here the results of the analyses performed when considering all the 210 cultivated accessions, the cluster cl5c being included. The inclusion of these accessions resulted in a lower fit to the two models considered here (*Supplementary Information*, Fig. S10-IIIA and S10-IVA) and in a shift of the inferred origin toward the West, in Central Africa, in a region encompassing the Southern Chad, the Western Sudan, the Central African Republic and the North of the Democratic Republic of Congo (*Supplementary Information*, Fig. S10-IIIB and C and S10-IVB and C). This shift may be due to the presence of accessions that are introgressed with wild sorghums and may reflect indeed a different evolutionary history for these sorghums from Western Africa.

#### *2.2 Estimation of the beginning of the expansion of the African cultivated sorghum and its spread rate*

We retrieved from the bibliography 15 African archeological sites with archaeobotanical remains of cultivated (or pre-domesticated) sorghum (Table S2-A) to calibrate our model of diffusion and estimate the timing of the onset of the expansion of the crop. The estimations considering all these sites led to a date for the latitudinal diffusion around 4,400 years ago (95% CI 4,167-4,791) (*Supplementary Information*, Fig. S11). The longitudinal diffusion was estimated to have started more recently, around 3,800 years ago (95% CI 3,641-4,224; *Supplementary Information*, Fig. S11). The longitudinal expansion was thus estimated to be more recent when the Ethiopian site was included. This is not surprising, as the Ethiopian site is geographically relatively close to the inferred origin of the expansion of the cultivated sorghum and is associated to relatively recent dates (300 BCE). This could represent the reality of a belated eastward spread or a bias due for example to the limited number of archeological sites in the area spanning older periods (Ruiz-Giralt et al., 2023) or to conservation issues impeding the retrieval of the oldest remains (Beldados and Ruiz-Giralt, 2023). The fact that most of our Ethiopian and easternmost accessions belonged to the more divergent and probably older clusters cl3 and cl4 would also argue in favor of the second hypothesis. In addition, for the domesticated sorghum to have been introduced in India by approx. 1,700 BCE from the Horn of Africa, at least pre-domesticated forms should have been already present in the area by that time (Ehret, 1979; Fuller and Stevens, 2018), making the hypothesis of a more recent eastward diffusion less plausible. Conversely, the estimations of the beginning of the latitudinal expansion with or without the Egyptian site were only slightly different (4,400 *vs.* 4,600 years ago). This could reflect the fact that the northward and the southward diffusions started approximately at the same period and/or that they dispersed at the same speed. More archeological sites North and East to the inferred origin of the expansion would be necessary to validate those results and hypotheses.

The estimation of the speed rate of diffusion considering all archeological sites led to a speed of 1.56 ± 0.81 km/year (80% CL: 0.34-2.73 km/year, r = 0.62) for the longitudinal and 1.22 ± 0.33 km/year (80% CL: 0.77-1.68 km/year, r = 0.82) for the latitudinal diffusion (*Supplementary Information*, Fig. S12-B).

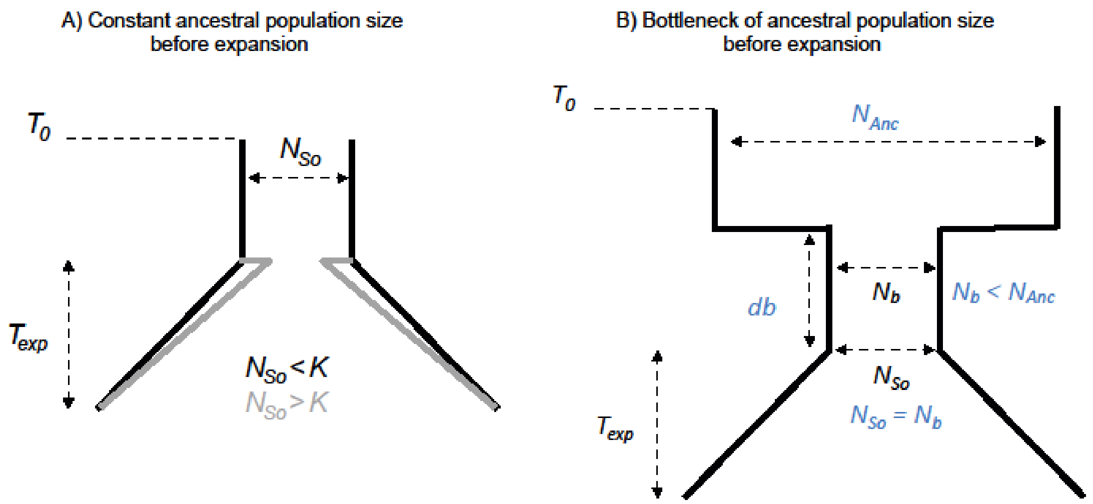


**Fig. S1**. Demographic scenarios of the cultivated sorghum in Africa simulated in this study. A) Model with a constant ancestral population size, of size $N_{So}$, before the expansion, which started $T_{exp}$ generation ago. $K$ is the carrying capacity defined as the number of individuals that can be sustained by the resources in a deme. When $N_{So}$, is greater than the carrying capacity, all individuals are found in the original deme, but the population of the source deme can be quickly downward regulated by the logistic growth function (grey lines representing this immediate bottleneck). B) Scenario involving a bottleneck of the ancestral population size before the expansion: before the expansion, which started $T_{exp}$ generation ago, the ancestral population (with a size of $N_{Anc}$) went through a bottleneck, decreasing the population size to $N_b$ during db generations. The size of the source population at the onset of the expansion ($N_{So}$) is equal to the size of the bottleneck populations ($N_b$).

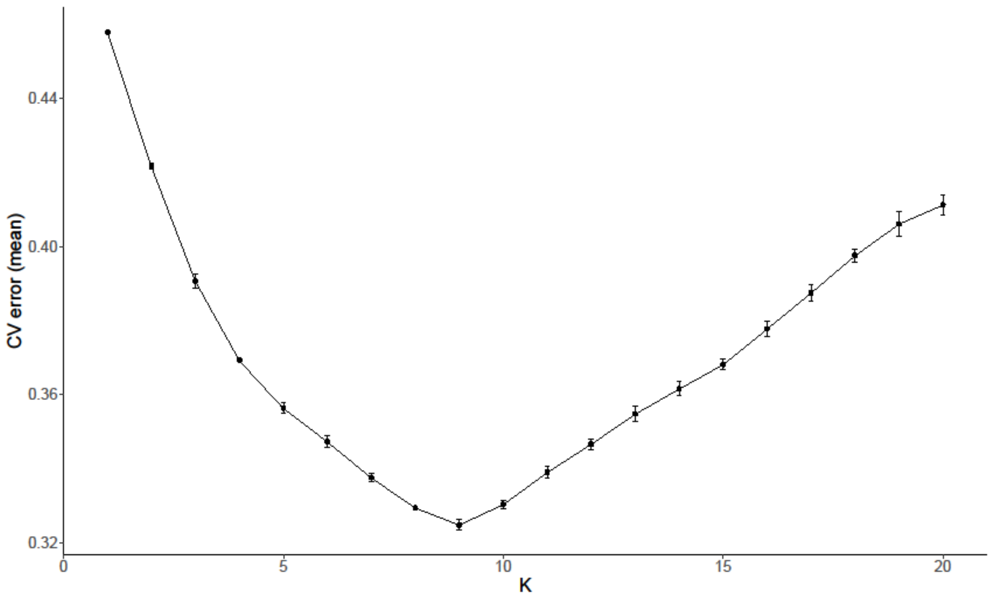


**Fig. S2.** Identification of the most plausible number of genetic groups in the panel of 210 African accessions of cultivated sorghum. The plot of the cross-validation error as a function of the number of genetic group *K* is given.

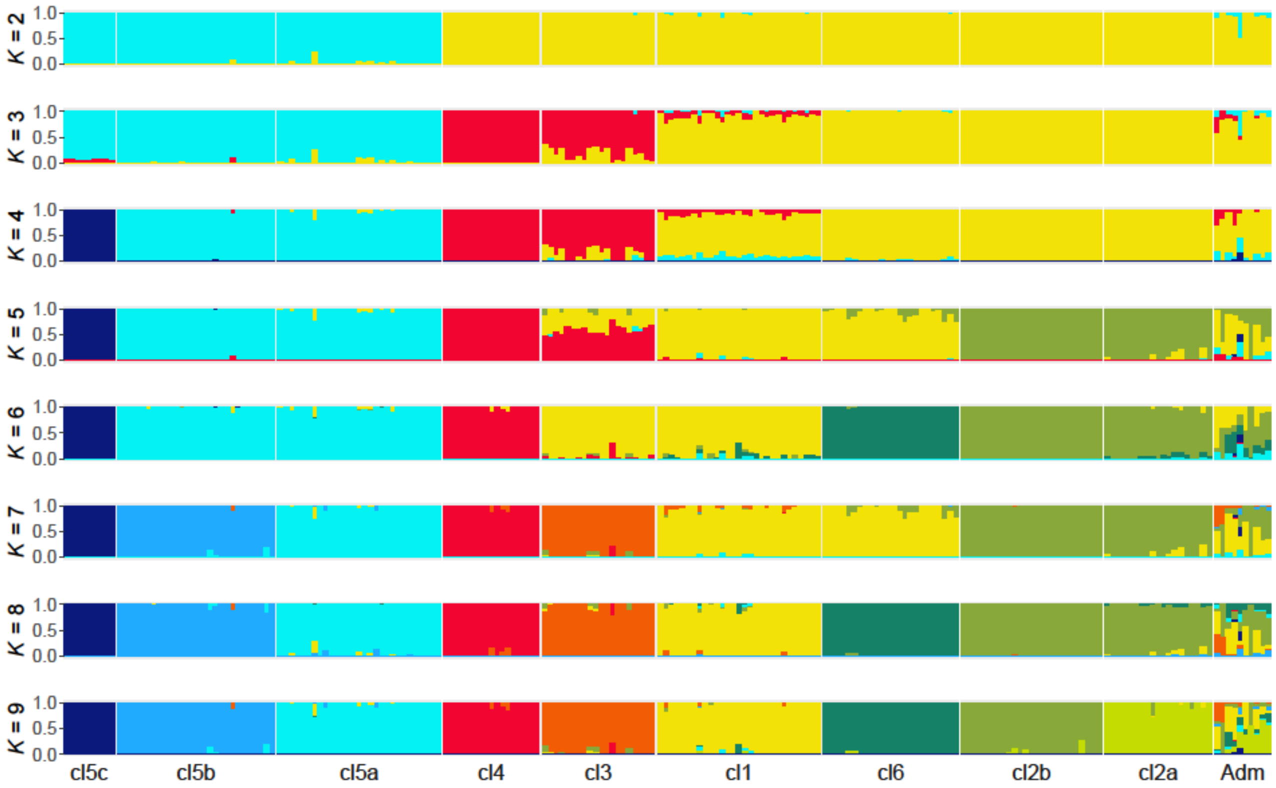


**Fig. S3.** Identification of the genetic clusters constituting our sampling when considering 2 (top) to 9 (bottom) genetic clusters and using the approach for model-based ancestry estimation implemented in ADMIXTURE (Alexander et al., 2009).

Each vertical line of the barplots corresponds to a sorghum accession, with the color representing the coefficient of ancestry for a genetic cluster. The accessions are arranged according to their geographic location, from West to South and then from North to South. Accessions that could not be assigned to a genetic cluster considering a threshold of 0.7 for the coefficient of ancestry are at the right extremity of the graph (Adm group).

A)

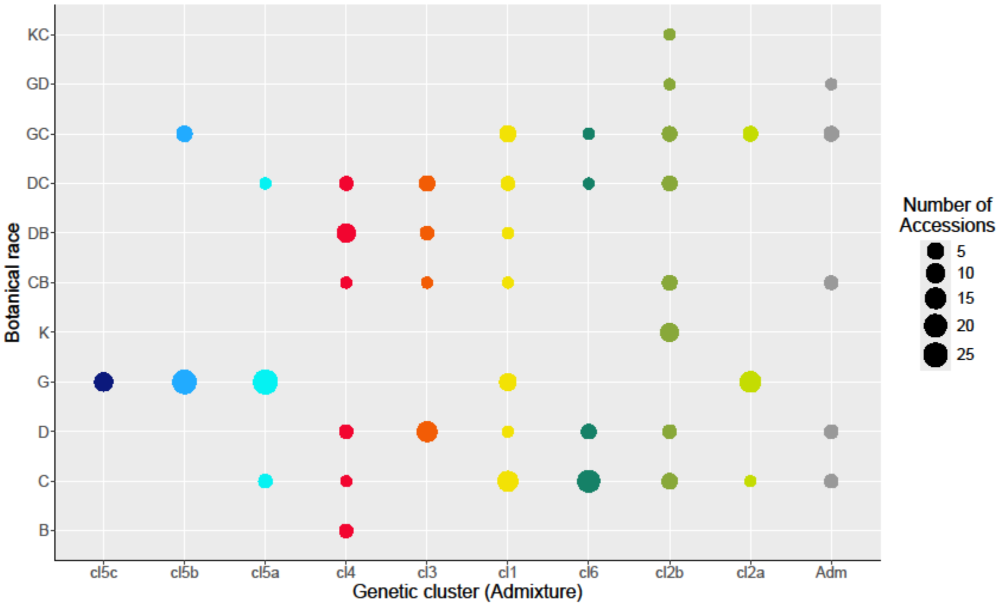


**B)**

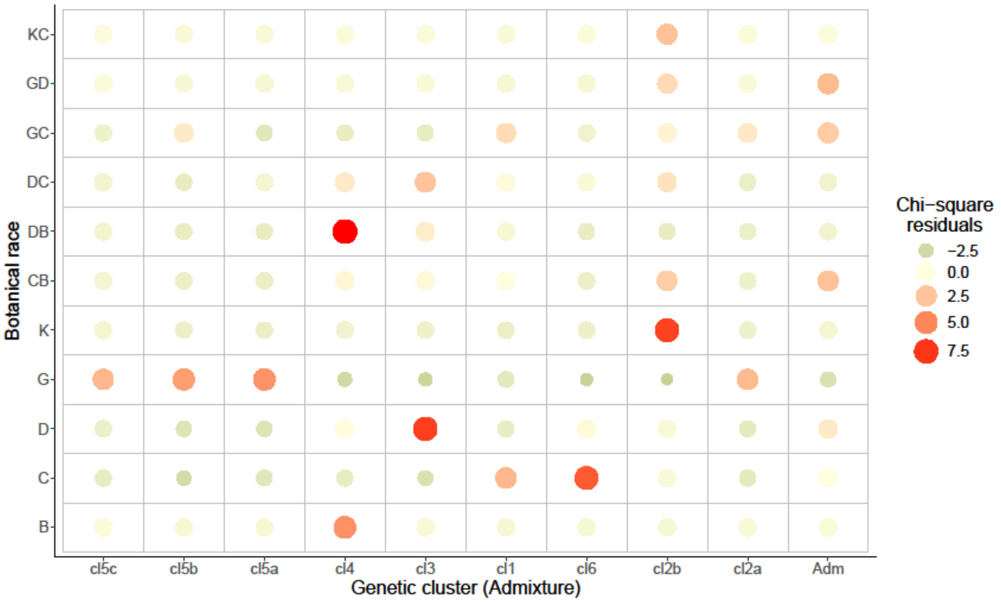


**Fig. S4.** A) Repartition of the sorghum accessions in the genetic clusters according to their botanical race. B) Plots of the Pearson residuals for the test of independence in the contingency table of the genetic clusters *vs.* the botanical races. B: bicolor; C: caudatum; D: durra; G: guinea; K: kafir. Genetic clusters are arranged according to their geographic location, from West to South and then from North to South.

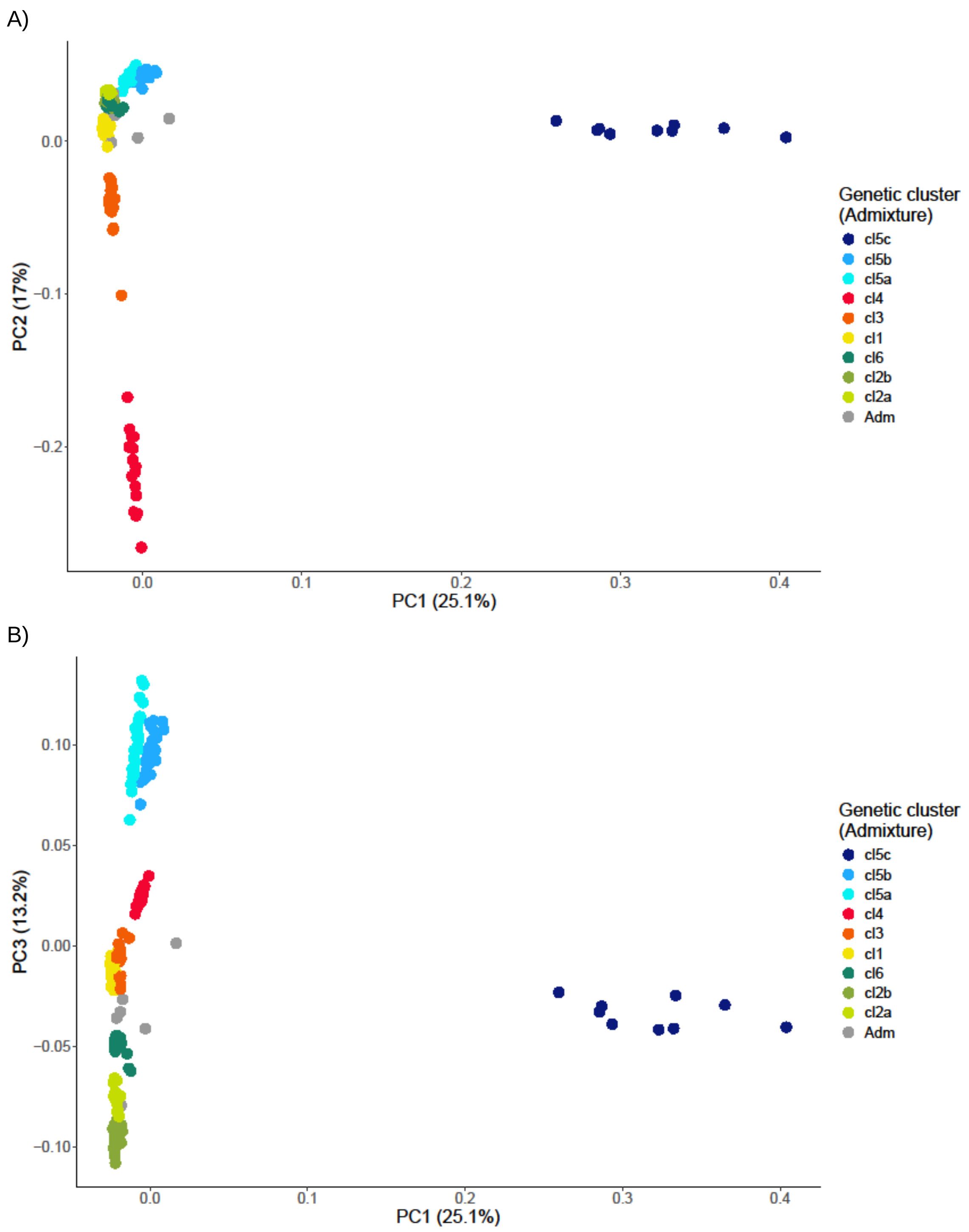
A)
PC2 (17%)
0.0
−0.1
−0.2
0.0
0.1
0.2
0.3
0.4
PC1 (25.1%)
Genetic cluster
(Admixture)
cl5c
cl5b
cl5a
cl4
cl3
cl1
cl6
cl2b
cl2a
Adm
B)
PC3 (13.2%)
0.10
0.05
0.00
−0.05
−0.10
0.0
0.1
0.2
0.3
0.4
PC1 (25.1%)
Genetic cluster
(Admixture)
cl5c
cl5b
cl5a
cl4
cl3
cl1
cl6
cl2b
cl2a
Adm

C)

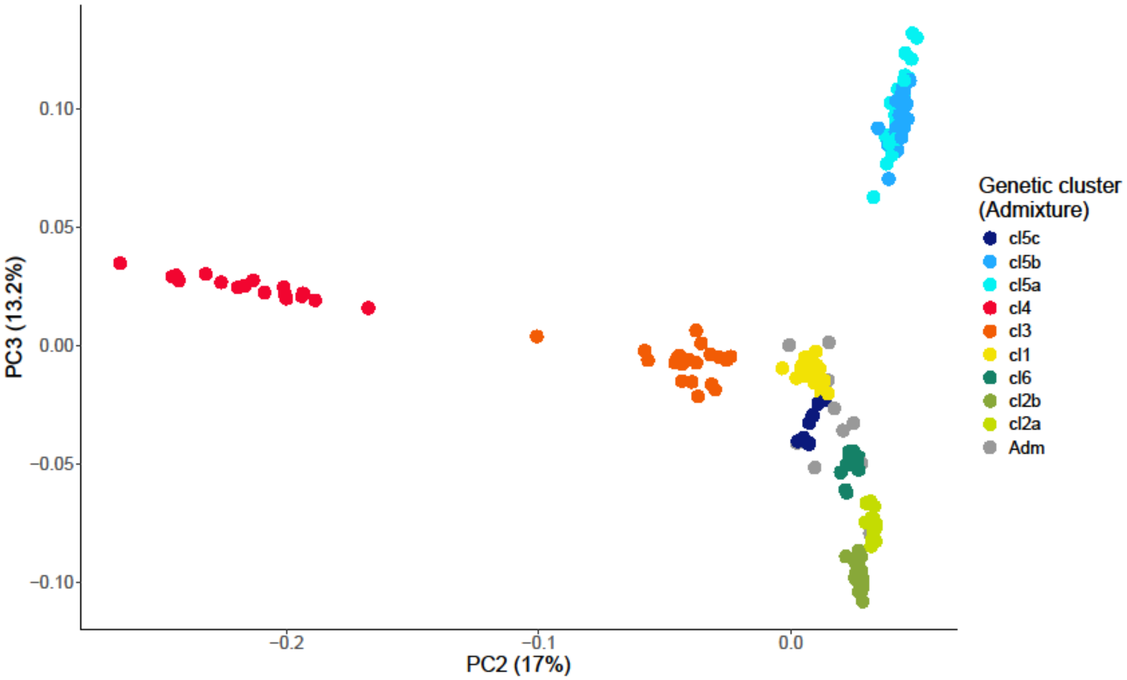


**Fig. S5.** Genetic diversity of the African cultivated sorghums assessed using a principal component analysis (PCA). A) Projection of the 210 sorghum accessions on the first and second axes that explain respectively 25.1 and 17% of the total variance. B) Projection of the sorghum accessions on the first and third axes that explain respectively 25.1 and 13.2% of the total variance. C) Projection of the accessions on the second (17% of the variance) and third axes (13.2%). Accessions are colored according to the ADMIXTURE clusters. Admixed accessions (gray color) are accessions that could not have been assigned unambiguously to a specific cluster, given a population ancestry threshold for assignation to a cluster of 0.7.

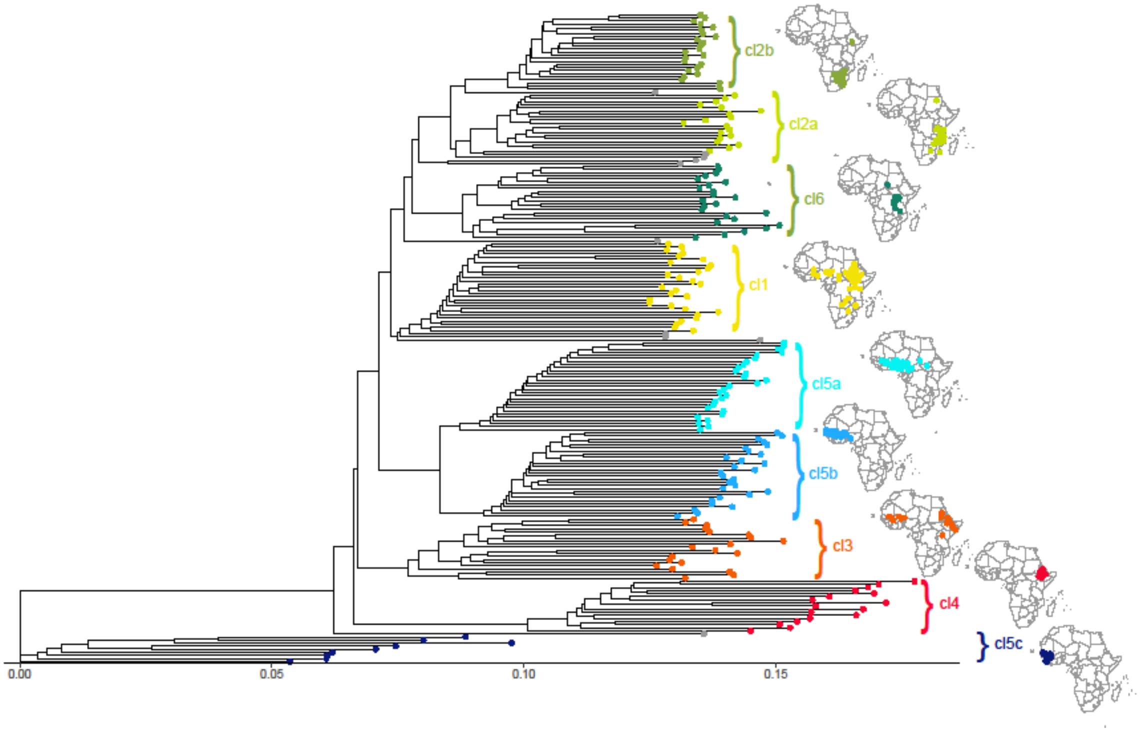


**Fig. S6.** Relationships of the 210 sorghum accessions represented using a Neighbor-joining tree. Accessions are colored according to the genetic groups defined using the clustering approach Admixture (Alexander et al., 2009). Admixed accessions (gray color) are accessions that could not be assigned unambiguously to a specific cluster, given a population ancestry threshold for assignation to a cluster of 0.7.

A)

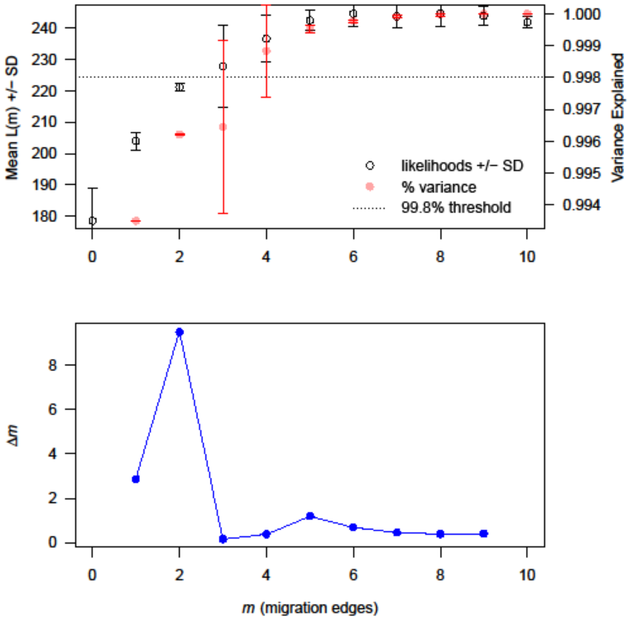


B)

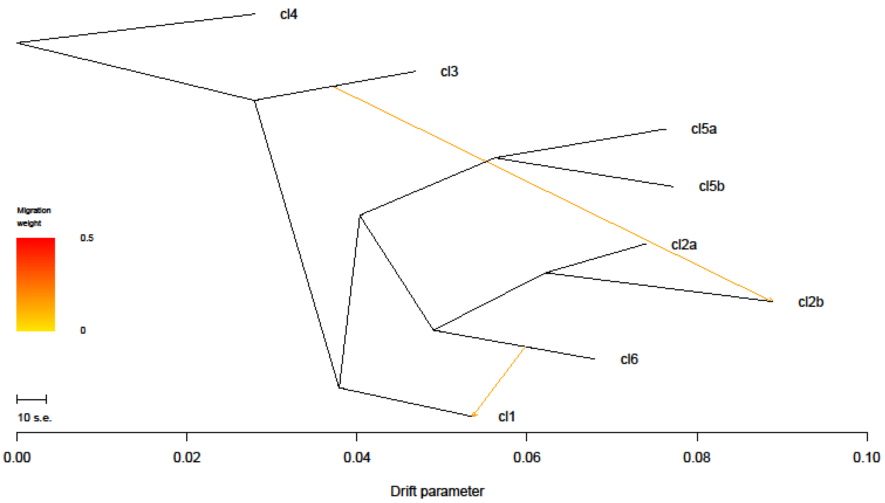


**Fig. S7.** TreeMix analysis of the relationships between the genetic clusters of the African cultivated sorghum. A) Estimation of the optimal number of migration events on the sorghum population tree using Treemix (Pickrell and Pritchard, 2012) and the R package OptM (Fitak, 2021). Top: Mean and standard deviation (SD) across 10 iterations for the composite likelihood L(m) and proportion of variance explained. The horizontal dotted line represented the 99.8% threshold recommended by Pickrell and Pritchard (2012). Bottom: The second-order rate of change (Δm) across values of m. B) Inferred tree with two migration events modeled

A)

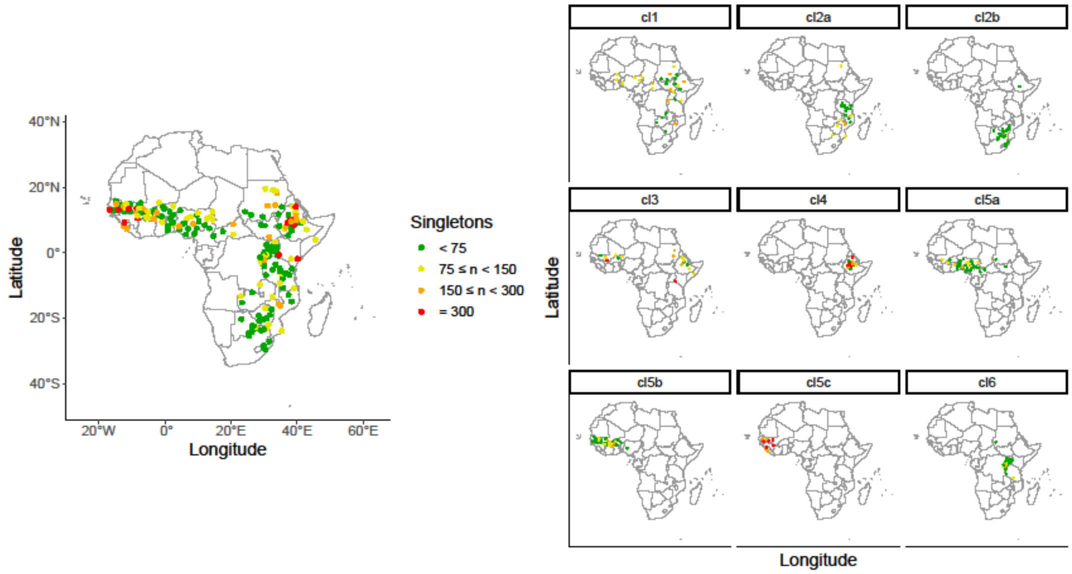


B)

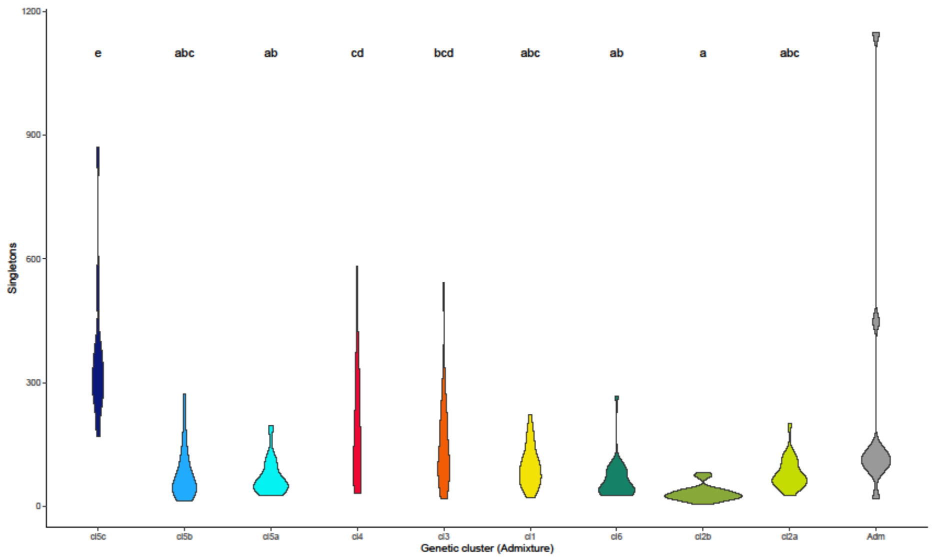


**Fig. S8.** Genetic diversity of the cultivated sorghum clusters represented using the number of singletons per accession. A) Geographic repartition of the singletons per accession, all accessions considered together (left) or for each cluster separately (right). B) Distribution of the number of singletons per accession within the genetic clusters. The letters at the top of the violin plots indicate the significance of Tukey's multiple comparisons test, with groups not differing in their mean of singletons sharing a letter.

A)

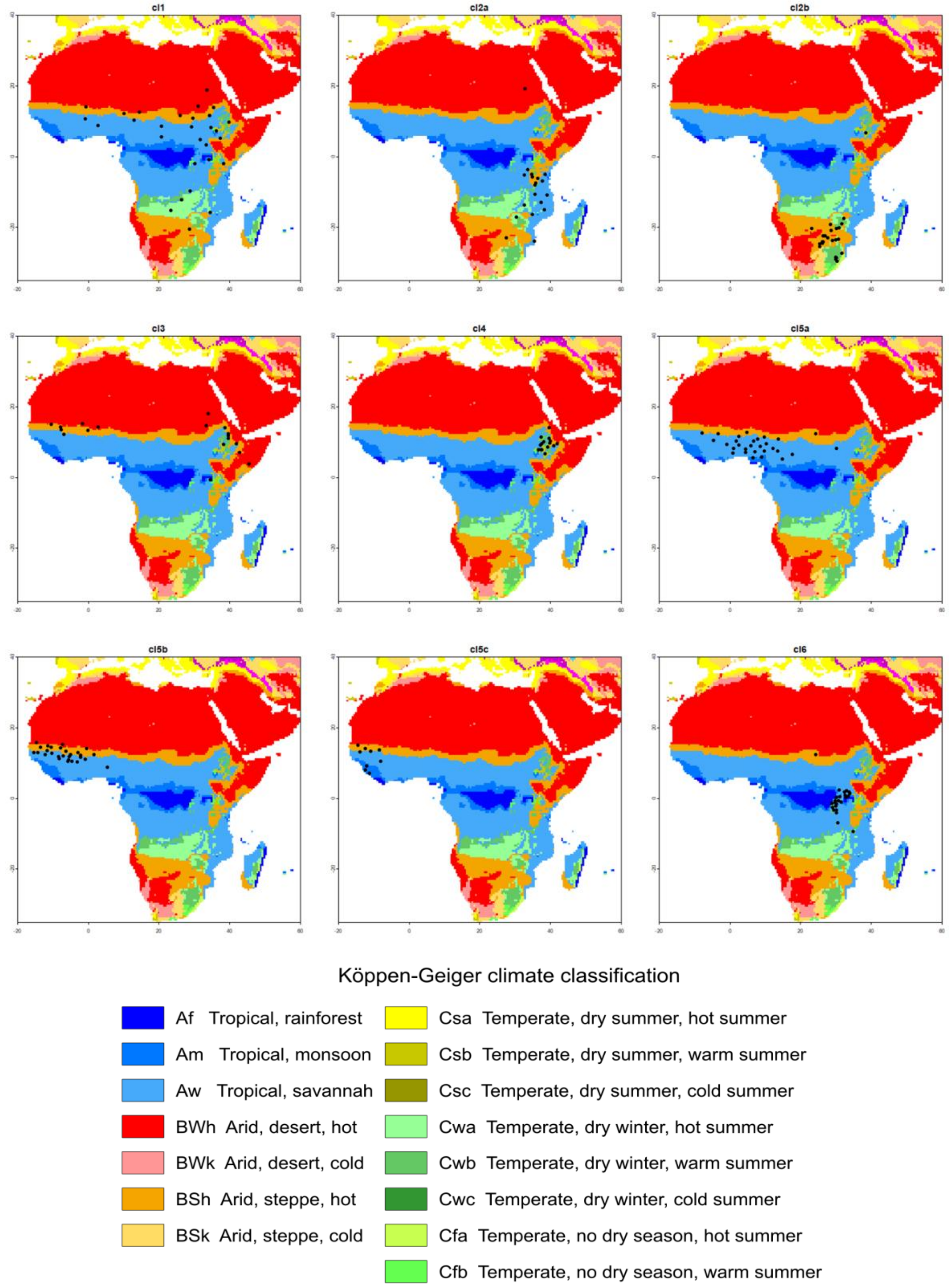
cl1
cl2a
cl2b
cl3
cl4
cl5a
cl5b
cl5c
cl6
Köppen-Geiger climate classification
Af Tropical, rainforest
Am Tropical, monsoon
Aw Tropical, savannah
BWh Arid, desert, hot
BWk Arid, desert, cold
BSh Arid, steppe, hot
BSk Arid, steppe, cold
Csa Temperate, dry summer, hot summer
Csb Temperate, dry summer, warm summer
Csc Temperate, dry summer, cold summer
Cwa Temperate, dry winter, hot summer
Cwb Temperate, dry winter, warm summer
Cwc Temperate, dry winter, cold summer
Cfa Temperate, no dry season, hot summer
Cfb Temperate, no dry season, warm summer

B)

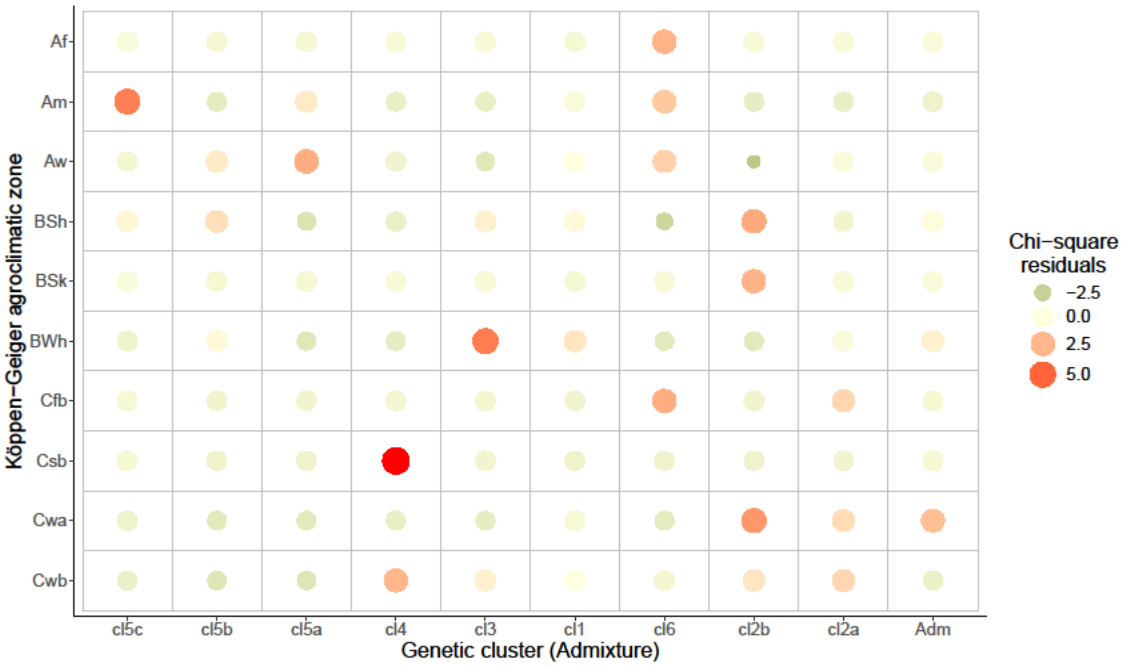
Af
Am
Aw
BSh
BSk
BWh
Cfb
Csb
Cwa
Cwb
Köppen−Geiger agroclimatic zone
cl5c
cl5b
cl5a
cl4
cl3
cl1
cl6
cl2b
cl2a
Adm
Genetic cluster (Admixture)
Chi−square residuals
−2.5
0.0
2.5
5.0

C)

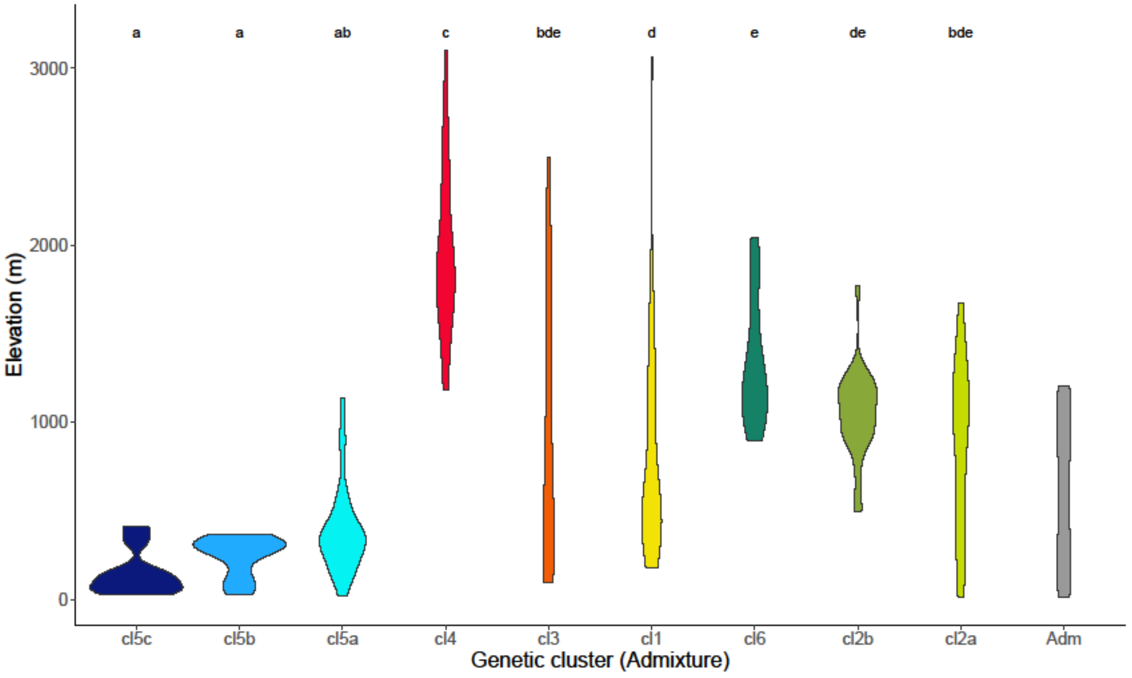
a
a
ab
c
bde
d
e
de
bde
3000
2000
1000
0
Elevation (m)
cl5c
cl5b
cl5a
cl4
cl3
cl1
cl6
cl2b
cl2a
Adm
Genetic cluster (Admixture)

D)

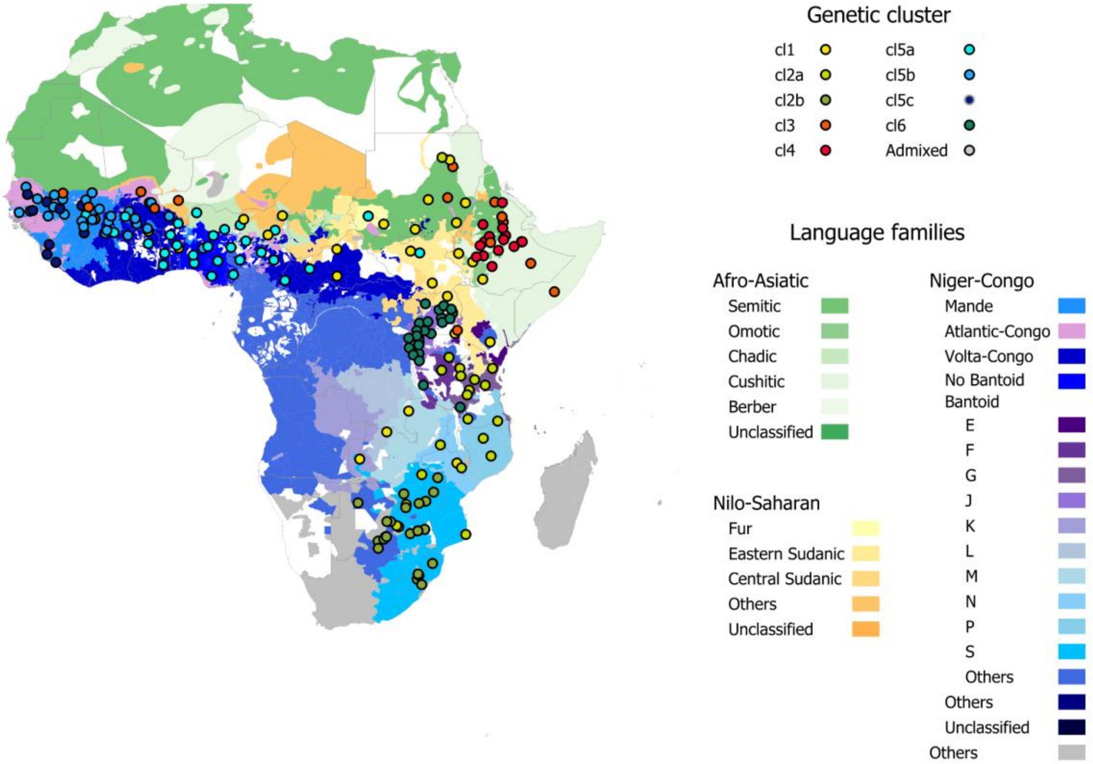


**Fig. S9.** Associations between genetic groups and elevation and agroclimatic or ethnolinguistic zones. A) Geographic distribution of the accessions according to the Köppen-Geiger agroclimatic classification. B) Plots of the Pearson residuals for the test of independence in the contingency table of the genetic clusters vs. the Köppen-Geiger agroclimatic zones. Circle size and color are proportional to the value of the residuals. C) Association between genetic clusters and the elevation. The letters at the top of the violin plots indicate the significance of Tukey's multiple comparisons test, with groups not differing in their mean elevation sharing a letter. D) Geographic distribution of the accessions according to the distribution of language families. Language data from Ethnologue version 16 (WLMS 16, www.gmi.org/wlms) (Lewis, 2009).

I) A)

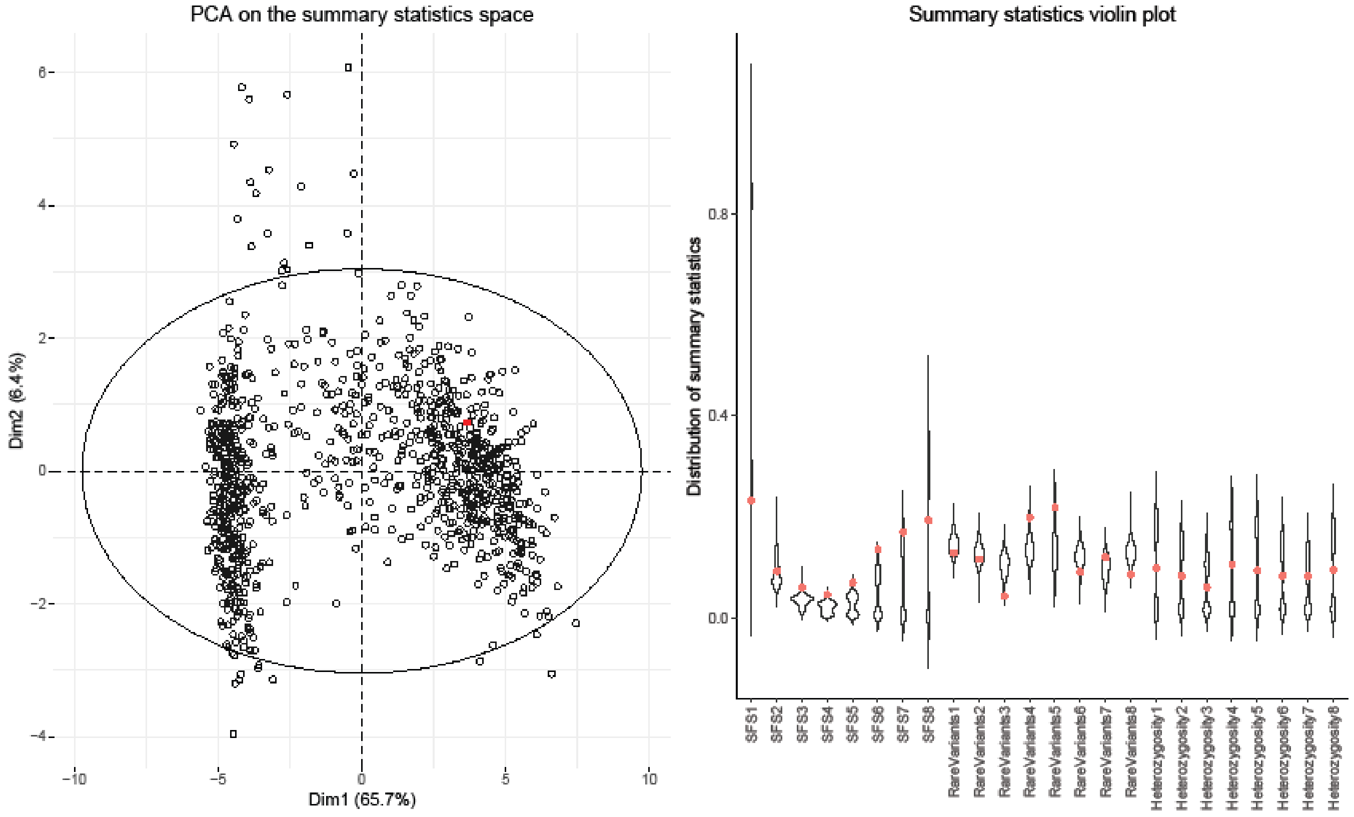
PCA on the summary statistics space
Dim1 (65.7%)
Dim2 (6.4%)
Summary statistics violin plot
Distribution of summary statistics
SFS1
SFS2
SFS3
SFS4
SFS5
SFS6
SFS7
SFS8
RareVariants1
RareVariants2
RareVariants3
RareVariants4
RareVariants5
RareVariants6
RareVariants7
RareVariants8
Heterozygosity1
Heterozygosity2
Heterozygosity3
Heterozygosity4
Heterozygosity5
Heterozygosity6
Heterozygosity7
Heterozygosity8

B)

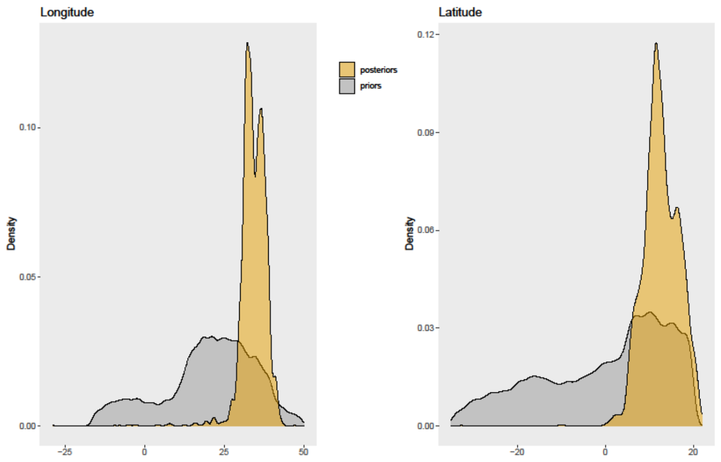
Longitude
posteriors
priors
Density
Latitude
Density

C)

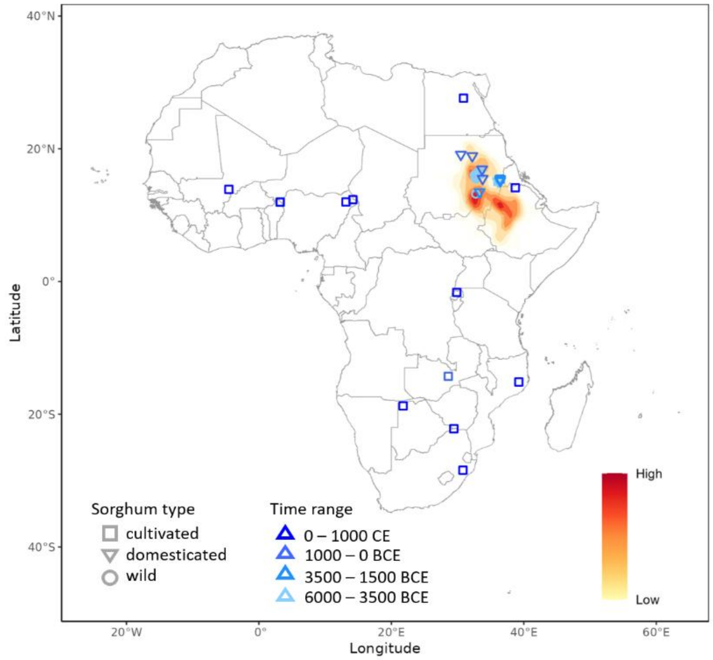
Latitude
Longitude
Sorghum type
cultivated
domesticated
wild
Time range
0 – 1000 CE
1000 – 0 BCE
3500 – 1500 BCE
6000 – 3500 BCE
High
Low

II) A)

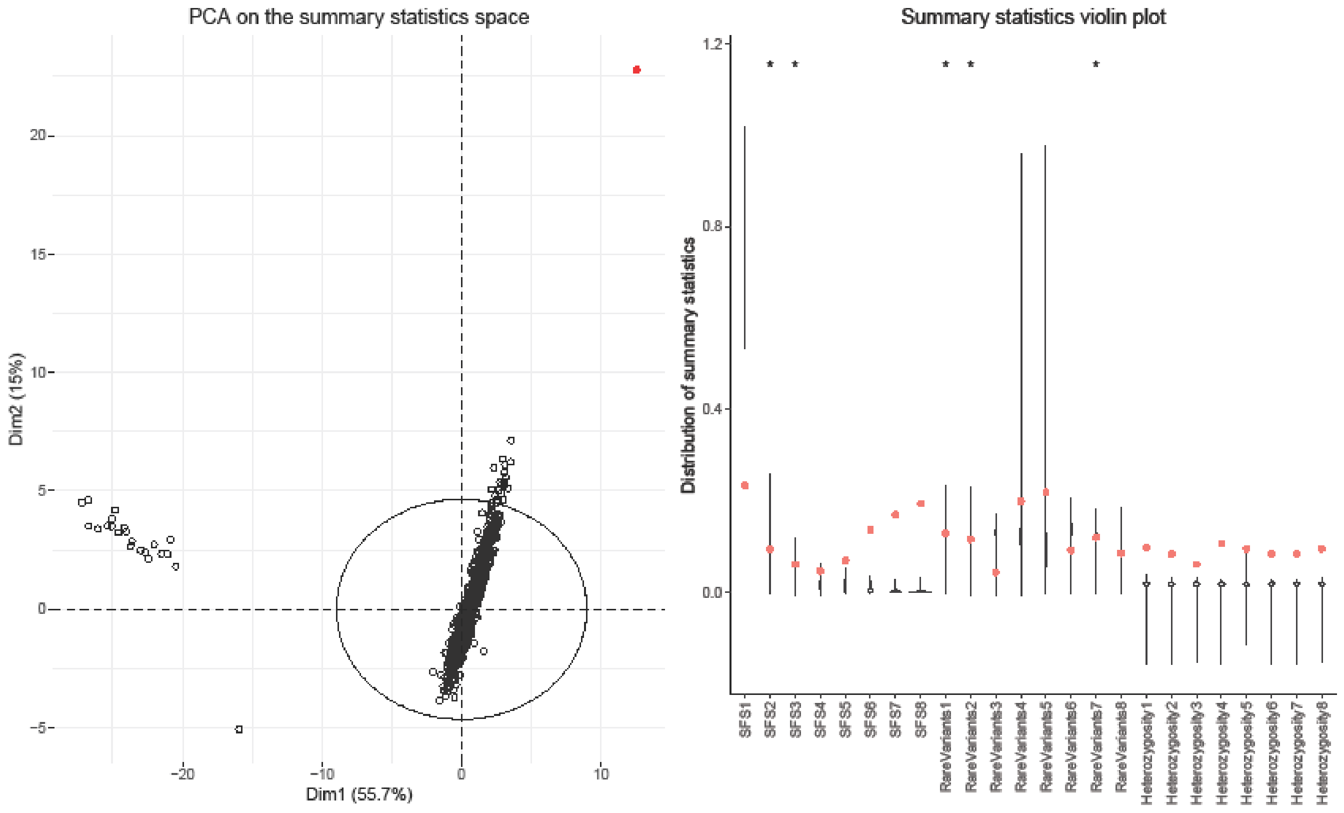
PCA on the summary statistics space
Dim1 (55.7%)
Dim2 (15%)
Summary statistics violin plot
Distribution of summary statistics
SFS1
SFS2
SFS3
SFS4
SFS5
SFS6
SFS7
SFS8
RareVariants1
RareVariants2
RareVariants3
RareVariants4
RareVariants5
RareVariants6
RareVariants7
RareVariants8
Heterozygosity1
Heterozygosity2
Heterozygosity3
Heterozygosity4
Heterozygosity5
Heterozygosity6
Heterozygosity7
Heterozygosity8

B)

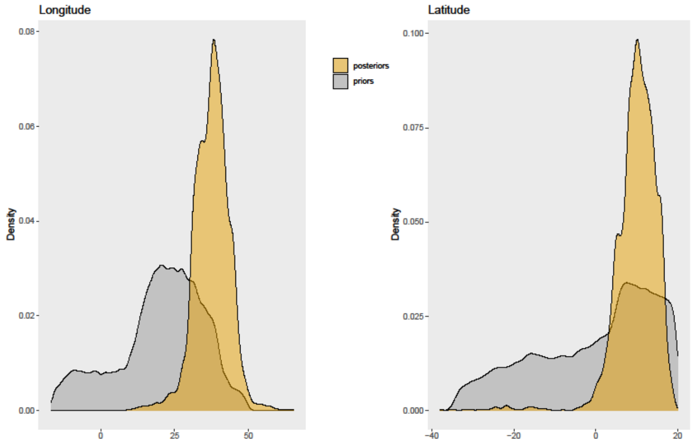
Longitude
Latitude
Density
posteriors
priors

C)

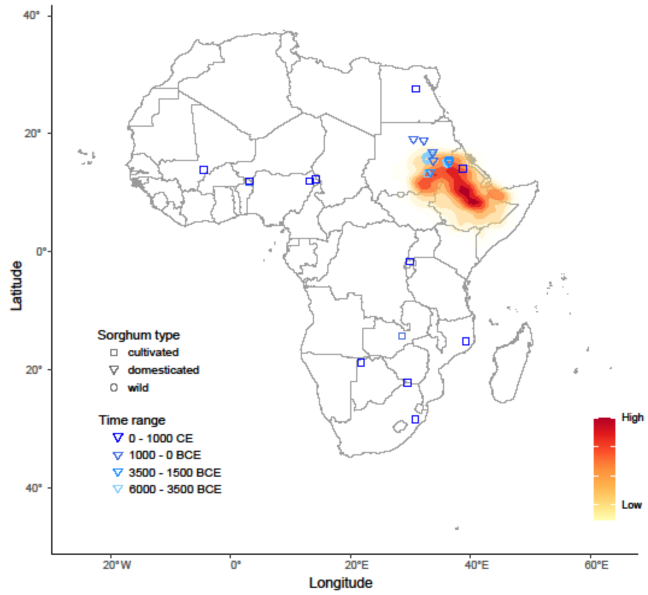
Latitude
Longitude
Sorghum type
cultivated
domesticated
wild
Time range
0 - 1000 CE
1000 - 0 BCE
3500 - 1500 BCE
6000 - 3500 BCE
High
Low

III) A)

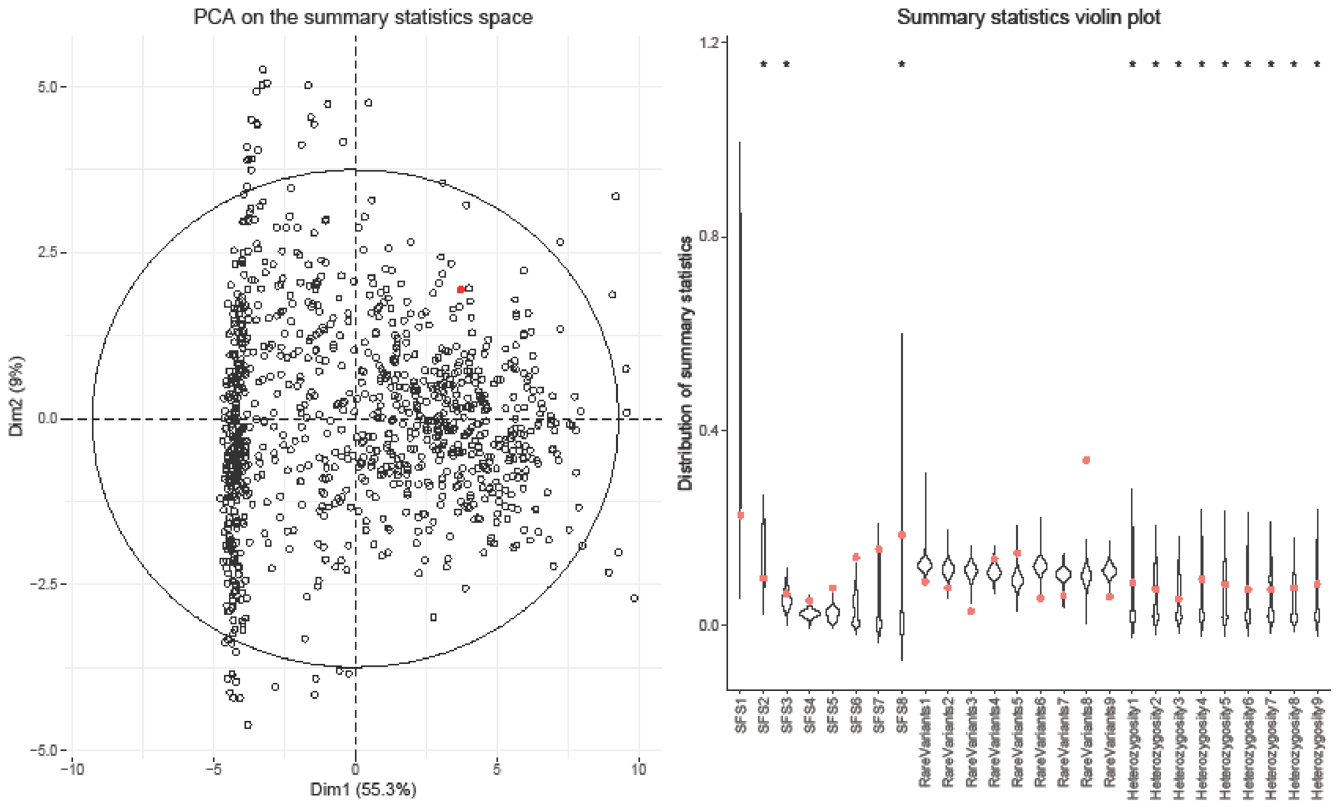

PCA on the summary statistics space
Dim2 (9%)
Dim1 (55.3%)
Summary statistics violin plot
Distribution of summary statistics
SFS1
SFS2
SFS3
SFS4
SFS5
SFS6
SFS7
SFS8
RareVariants1
RareVariants2
RareVariants3
RareVariants4
RareVariants5
RareVariants6
RareVariants7
RareVariants8
RareVariants9
Heterozygosity1
Heterozygosity2
Heterozygosity3
Heterozygosity4
Heterozygosity5
Heterozygosity6
Heterozygosity7
Heterozygosity8
Heterozygosity9


B)

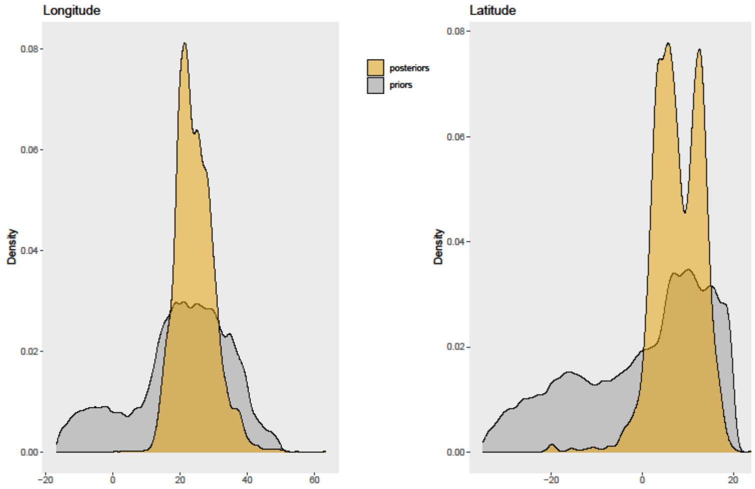

Longitude
Latitude
Density
posteriors
priors


C)

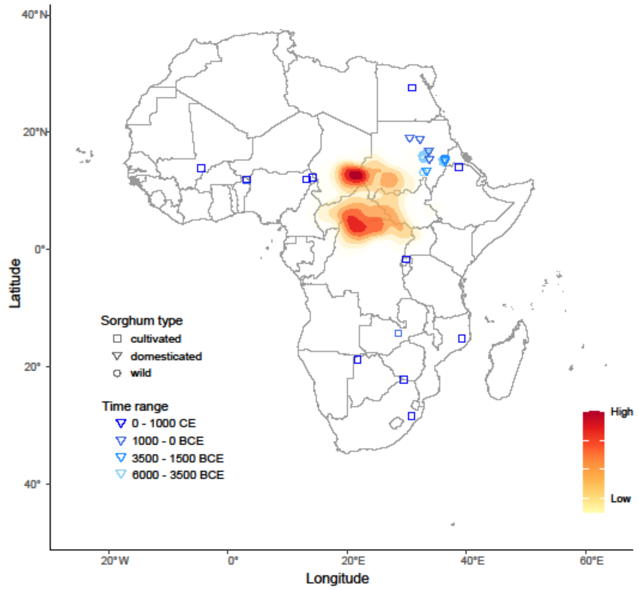

Latitude
Longitude
Sorghum type
cultivated
domesticated
wild
Time range
0 - 1000 CE
1000 - 0 BCE
3500 - 1500 BCE
6000 - 3500 BCE
High
Low

IV) A)

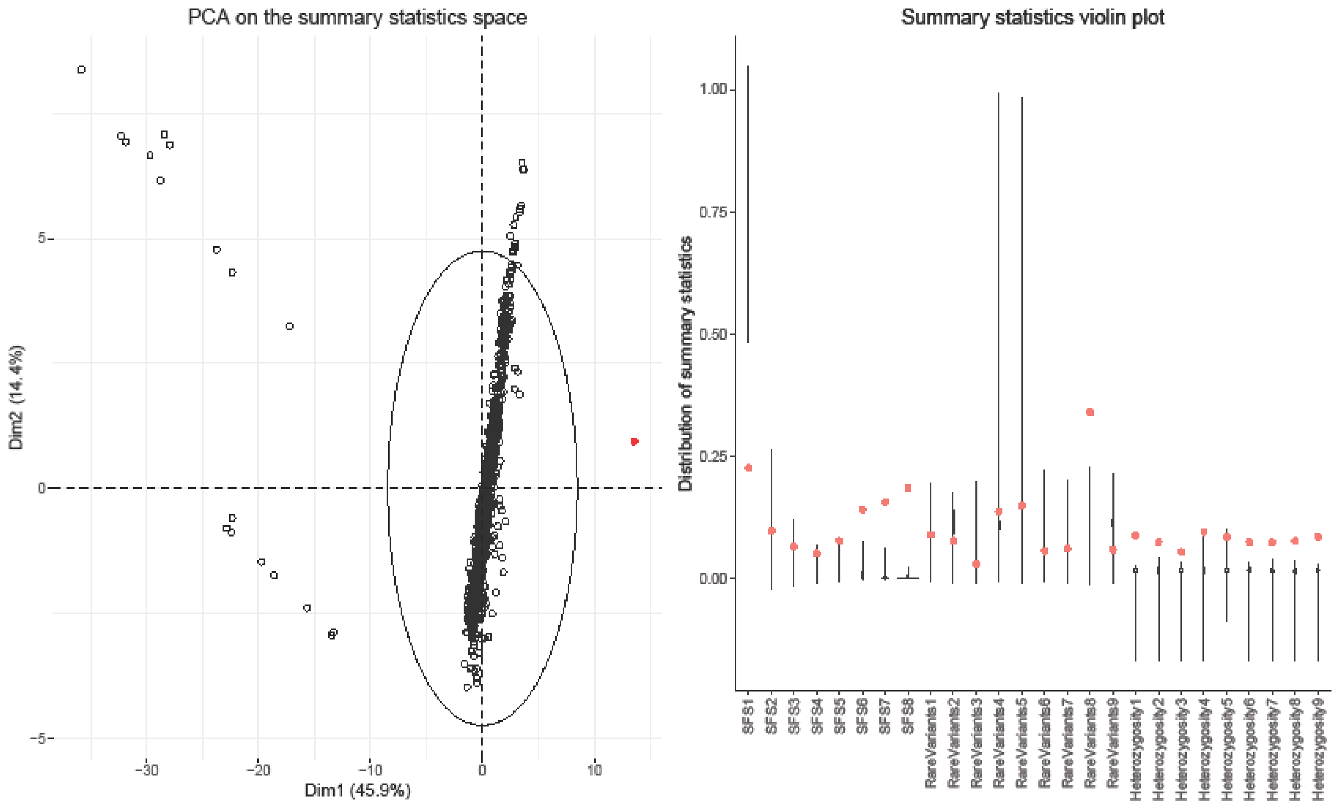
PCA on the summary statistics space
Dim1 (45.9%)
Dim2 (14.4%)
Summary statistics violin plot
Distribution of summary statistics
SFS1
SFS2
SFS3
SFS4
SFS5
SFS6
SFS7
SFS8
RareVariants1
RareVariants2
RareVariants3
RareVariants4
RareVariants5
RareVariants6
RareVariants7
RareVariants8
RareVariants9
Heterozygosity1
Heterozygosity2
Heterozygosity3
Heterozygosity4
Heterozygosity5
Heterozygosity6
Heterozygosity7
Heterozygosity8
Heterozygosity9

B)

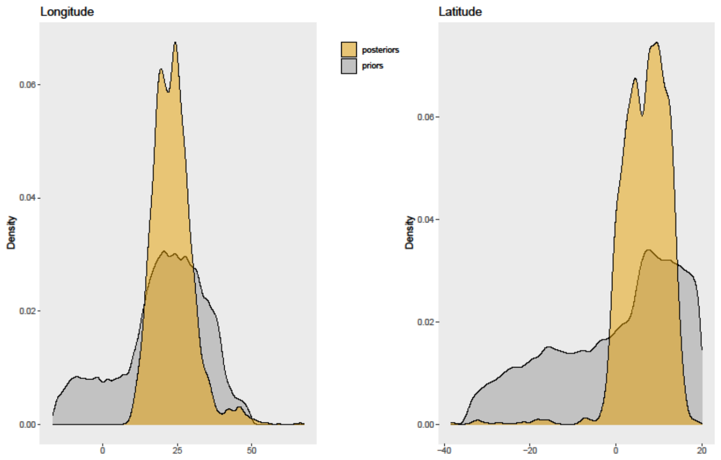
Longitude
Latitude
Density
posteriors
priors

C)

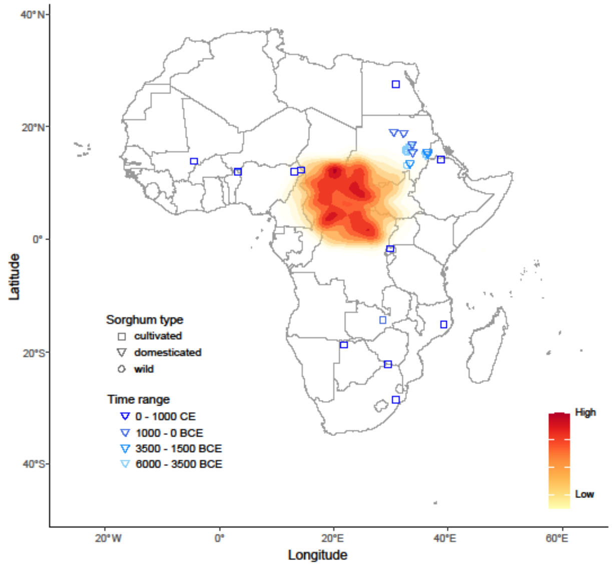
Latitude
Longitude
Sorghum type
cultivated
domesticated
wild
Time range
0 - 1000 CE
1000 - 0 BCE
3500 - 1500 BCE
6000 - 3500 BCE
High
Low

**Fig. S10.** ABC analyses of the geographic origin of the cultivated sorghum in Africa. ABC analyses on the dataset of all the cultivated accessions but the cluster cl5c (I and II) and on the complete dataset of cultivated sorghums (III and IV), considering either a model of constant population size before the expansions (I and III) or a model of ancestral population bottleneck before the expansions of cultivated sorghum in Africa (II and IV). For each analysis, we provide: (A) the posterior predictive checks for the model. We provide on the left the PCA on the summary statistics and on the right the violin plots representing the distributions of the summary statistics calculated from 1000 simulated dataset made using parameters drawn from the posterior distributions resulting from the ABC analysis. The red dots represent the observed values on the real dataset, and the ellipse on the PCA shows the 95% confidence ellipse); (B) the prior and posterior distributions for the latitude and longitude; and (C) the map of the inferred origin with the projection of 2D-kernel density estimates. The blue symbols correspond to the localizations of the archeological sites in the inferred area of origin where the oldest remains of sorghum were found (from Table S4-B).

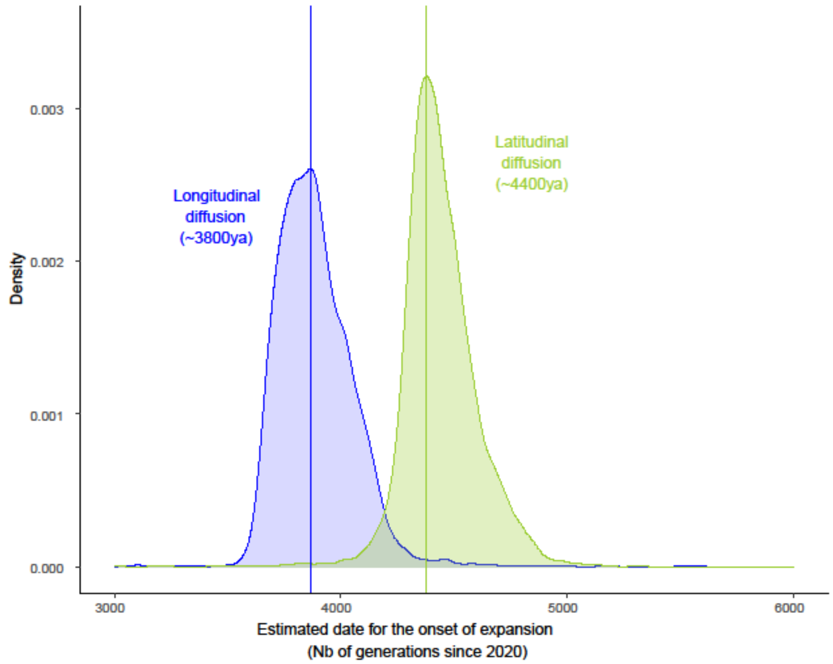


**Fig. S11.** Distribution of the timing of the beginning of the longitudinal (blue) and latitudinal (green) movements of diffusion of the African cultivated sorghum, considering the 15 archeological sites.

A)

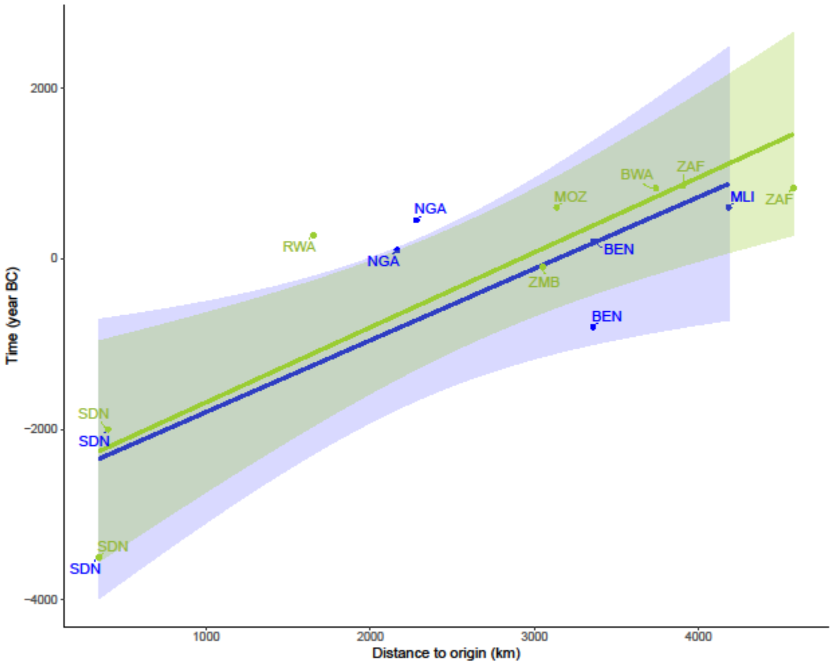


B)

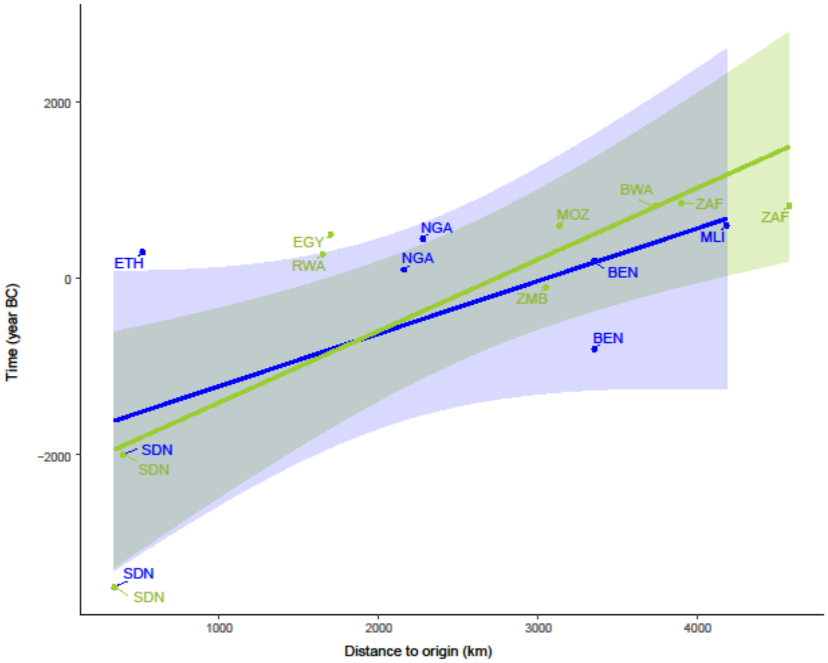


**Fig. S12**. Estimation of the speed of the fronts of diffusion of cultivated sorghum in Africa. A) Linear regression fit to estimate the speed of the southward and westward fronts of dispersal of the cultivated sorghum in Africa. The Egyptian and Ethiopian sites were not considered here to fit the regressions. B) Linear regression fit to estimate the speed of the fronts of dispersal of the cultivated sorghum in Africa using the 15 archeological sites considered in this study. Blue: westward expansion, green: southward expansion.

**Table S1.** Splatche parameters.
A) Priors' distributions of the model parameters

| Parameter name | | Acronym | Parameter description | Distribution | Constant ancestral population size | | Bottleneck before expansion | |
|---|---|---|---|---|---|---|---|---|
| | | | | | min | max | min | max |
| AncSize | | Nanc | size of the ancestral population (nb of haploid individuals) / population size before the bottleneck | uniform | 0 | | 300 | 100000 |
| SoSize | | NSo | size of the source population | uniform | 300 | 100000 | =Nb | |
| SoLong | | LgSo | Longitude of the source population | uniform | -17 | 50 | | |
| SoLat | | LtSo | Latitude of the source population | uniform | -35 | 20 | | |
| CarrCap | | K | Carrying capacity | uniform | 10 | 150 | | |
| EndTime | | Texp | Number of simulated generations | uniform | 100 | 3000 | | |
| GrowthRate | | r | Growth rate | uniform | 0,05 | 1 | | |
| MigrationRate | | m | Migration rate for neighboring deme migration | uniform | 0,05 | 0,9 | | |
| BtSize | | Nb | Size of the source population before the start of the expansion & after the bottleneck (if set to 0, the size of the source pop regarded as being equal to the initial size set in the 2nd col) | uniform | NA | | 100 | min(3000, AncSize) |
| BtDuration | | db | ~duration of the bottleneck | uniform | NA | | 100 | 3000 |

B) Fixed parameters

| Parameter name | Acronym | Parameter description | Value |
|---|---|---|---|
| Friction | F | friction | 0,5 |
| GenerationTime | G | Duration of a generation, in years | 1 |
| realBPTime | T0 | Real time, in years before present, of the start of the simulation | -4000 |
| RiverFrictionChangeFactor | | Factor that increases or decreases the friction over the river cells | 0,5 |
| RiverCarCapChangeFactor | | Factor that increases or decreases the carrying cap over the river cells | 2 |
| CoastFrictionChangeFactor | | Factor that increases or decreases the friction over the coast cells | 0,5 |
| CoastCarCapChangeFactor | | Factor that increases or decreases the carrying cap over the coast cells | 2 |

C) Model choice parameters

| C) Model choice parameters | Parameter description | Constant ancestral population size | Bottleneck before expansion |
|---|---|---|---|
| DemographicModel | demographic model | 3 | |
| AllowSourcePopOverflow | density N attributed to the initial deme spread over neighboring demes if N greater than K | 1 | 0 |
| FrictionChoice | vegetation + roughness topo | 2 | |
| NumGeneticSimulations | number of genetic simulations following the demographic simulation | 10 | |
| GenotypicData | 1: genotypic / 0: haplotypic | 1 | |
| MaxNumGenerations | maximum number of total generations for a genetic simulation | 1000000 | |

**Table S2.** Archeological sites with known (pre-)domesticated sorghum remains.
A) Known archeological sites throughout the African area of distribution of sorghum. Long. and Lat. diffusion stand respectively for longitudinal and latitudinal diffusion.

| **Site** | **Main Region** | **Country** | **Latitude** | **Longitude** | **Earliest Date BC/AD** | **Long. diffusion** | **Lat. diffusion** | **References** |
|---|---|---|---|---|---|---|---|---|
| Khashm el Girba (KG23) | Africa NE | Sudan | 15.076886 | 36.101654 | -3500 | | | Barron et al., 2020 |
| Mahal Teglinos (K1) | Africa NE | Sudan | 15.442182 | 36.431245 | -2000 | | | Beldados et al., 2018 |
| Alibori SIII | Africa West | Benin | 11.945200 | 3.220000 | -800 | | | Champion and Fuller, 2018 |
| M'teteshi | Africa South | Zambia | -14.279170 | 28.595830 | -100 | | | Robertson, 1991 |
| Kursakata | Africa West | Nigeria | 12.327248 | 14.202014 | 100 | | | Zach and Klee, 2003 |
| Birni Lafia | Africa West | Benin | 11.966450 | 3.221000 | 200 | | | Champion and Fuller, 2018 |
| Kabusanze (BPS36) | Africa East | Rwanda | -1.640278 | 29.878889 | 275 | | | Giblin and Fuller, 2011 |
| Axum | Africa NE | Ethiopia | 14.131942 | 38.719491 | 300 | | | Boardman, 2000, 1999 |
| Dorota | Africa West | Nigeria | 11.982183 | 13.141878 | 450 | | | Magnavita, 2002 |
| Kom el-Nana | Africa NE | Egypt | 27.619404 | 30.889040 | 500 | | | Smith, 2003 |
| Jenne-Jeno | Africa West | Mali | 13.878494 | -4.539905 | 600 | | | Mcintosh, 1995 |
| Xakota (Nampula) | Africa South | Mozambique | -15.14000 | 39.250000 | 600 | | | Boivin et al., 2013; Sinclair et al., 1993 |
| Magogo | Africa South | South Africa | -28.43361 | 30.806670 | 825 | | | Maggs and Ward, 1984; Mitchell, 2002 |
| Nqoma | Africa South | Botswana | -18.75000 | 21.766670 | 825 | | | Denbow and Wilmsen, 1986; Mitchell, 2002 |
| Schroda | Africa South | South Africa | -22.17667 | 29.433610 | 850 | | | Hanisch, 1981; Mitchell, 2002 |

B) Earliest archeological sites with known (pre-)domesticated sorghum remains in Sudan, the inferred area of origin

| Site | Latitude | Longitude | Date (BCE) | Sorghum type | References |
|---|---|---|---|---|---|
| Sheikh el-Amin | 15.579444 | 32.8275 | 6000-3500 | Wild | Abdel-Magid, 2003 |
| El Mahalab | 15.603333 | 32.8066667 | 6000-3500 | Wild | Abdel-Magid, 2003 |
| Ghaba | 16.520017 | 33.126894 | 6000-3500 | Wild | Madella et al., 2014 |
| Um Direiwa | 15.721685 | 32.661236 | 6000-3500 | Wild | Abdel-Magid, 1989; Stemler, 1990 |
| Ghaba | 16.520017 | 33.126894 | 6000-3500 | Wild | Madella et al., 2014 |
| Rabak | 13.181067 | 32.7424303 | 6000-3500 | Wild | Abdel-Magid, 1989 |
| Sheikh Mustafa | 15.483333 | 32.7666667 | 6000-3500 | Wild | Abdel-Magid, 2003 |
| El Zakiab | 15.773752 | 32.57755 | 6000-3500 | Wild | Abdel-Magid, 1989; Stemler, 1990 |
| Shaheinab | 16.06977 | 32.540777 | 6000-3500 | Wild | Abdel-Magid, 1989 |
| El Kadada | 16.804669 | 33.532216 | 6000-3500 | Wild | Stemler, 1990 |
| Kadero | 15.757974 | 32.609018 | 6000-3500 | Wild | Stemler, 1990 |
| Khashm el Girba (KG23) | 15.076886 | 36.101654 | 6000-3500 | Domestic | Winchell et al., 2017 |
| Shaqadud cave | 16.230003 | 33.38213 | 6000-3500 | Cultivated | Abdel-Magid, 1989 |
| Jebel Moya | 13.48442 | 33.31936 | 3500-1500 | Domestic | Brass et al., 2019 |
| Mahal Teglinos (K1) | 15.442182 | 36.431245 | 3500-1500 | Domestic | Beldados et al., 2018 |
| Kassala, JAG 9/SEG 1 | 15.170289 | 36.435361 | 3500-1500 | Domestic | Costantini et al., 1983, 1982 |
| Kawa | 19.126537 | 30.496467 | 1000-0 | Domestic | D. Q. Fuller, 2004 |
| Umm Muri | 18.914427 | 32.218034 | 1000-0 | Domestic | D. Fuller, 2004; Fuller, 2014 |
| Hamadab | 16.914765 | 33.693985 | 1000-0 | Domestic | Fuller, et al, N.D. |
| Jebel Qeili | 15.507115 | 33.784933 | 1000-0 | Domestic | Fuller, 2014 |

**Table S3.** Genetic diversity of the 9 genetic clusters identified within the panel of 210 cultivated African sorghums analyzed in this study.

N: number of accessions; pA: number of private alleles; pAn: normalized number of private alleles; singletons: mean number of singletons per accession; $H_O$: the observed heterozygosity; $H_S$: the gene diversity; $F_{IS}$: Wright's fixation index.

| Genetic cluster | N | *pA* | *pAn* | singletons | *Ho* | *Hs* | *Fis* | *Fis* 95% CI |
|---|---|---|---|---|---|---|---|---|
| cl1 | 29 | 4273 | 147.3 | 97.4 | 0.0099 | 0.087 | 0.8864 | [0.8428-0.8609] |
| cl2a | 19 | 2373 | 124.9 | 80.5 | 0.0116 | 0.0762 | 0.848 | [0.7809-0.801] |
| cl2b | 25 | 1358 | 54.3 | 31.8 | 0.0065 | 0.0550 | 0.8824 | [0.8278-0.8535] |
| cl3 | 20 | 4929 | 246.5 | 156 | 0.0121 | 0.0965 | 0.8746 | [0.84-0.8555] |
| cl4 | 17 | 6751 | 397.1 | 182 | 0.0169 | 0.0874 | 0.8062 | [0.7628-0.784] |
| cl5a | 29 | 3572 | 123.1 | 70.2 | 0.0144 | 0.0742 | 0.8052 | [0.7191-0.7417] |
| cl5b | 28 | 3163 | 113 | 81.5 | 0.0175 | 0.0737 | 0.7621 | [0.6975-0.7199] |
| cl5c | 9 | 8822 | 980.2 | 385 | 0.02 | 0.0912 | 0.7803 | [0.7026-0.7249] |
| cl6 | 24 | 3153 | 131.4 | 64.5 | 0.0097 | 0.0854 | 0.8859 | [0.8396-0.8586] |

**Table S4.** Distribution of the cultivated accessions of each sorghum cluster according to the Geiger-Köpper climate zone (A) or the language family of their location (B). The four classes of family language considered at the higher level are highlighted in blue.

A)

| Clusters | Af Tropical, rainforest | Am Tropical, monsoon | Aw Tropical, savanna | BSh Arid, steppe, hot | BSk Arid, steppe, cold | BWh Arid, desert, hot | Cfb Temperate, no dry season, warm summer | Csb Temperate, dry summer, warm summer | Cwa Temperate, dry winter, hot summer | Cwb Temperate, dry winter, warm summer |
|---|---|---|---|---|---|---|---|---|---|---|
| cl1 | 0 | 1 | 13 | 9 | 0 | 3 | 0 | 0 | 1 | 2 |
| cl2a | 0 | 0 | 8 | 4 | 0 | 1 | 1 | 0 | 2 | 3 |
| cl2b | 0 | 0 | 1 | 15 | 1 | 0 | 0 | 0 | 5 | 3 |
| cl3 | 0 | 0 | 5 | 7 | 0 | 6 | 0 | 0 | 0 | 2 |
| cl4 | 0 | 0 | 6 | 3 | 0 | 0 | 0 | 4 | 0 | 4 |
| cl5a | 0 | 2 | 23 | 4 | 0 | 0 | 0 | 0 | 0 | 0 |
| cl5b | 0 | 0 | 15 | 11 | 0 | 2 | 0 | 0 | 0 | 0 |
| cl5c | 0 | 3 | 3 | 3 | 0 | 0 | 0 | 0 | 0 | 0 |
| cl6 | 1 | 3 | 16 | 1 | 0 | 0 | 2 | 0 | 0 | 1 |
| Admixed | 0 | 0 | 4 | 3 | 0 | 1 | 0 | 0 | 2 | 0 |

B)

| | | | Niger-Congo | | | | | | | | | | | | | | | | | |
|---|---|---|---|---|---|---|---|---|---|---|---|---|---|---|---|---|---|---|---|---|
| | | | | Atlantic-Congo | | | | | | | | | | | | | | | | |
| | | | | | | Volta-Congo | | | | | | | | | | | | | | |
| | | | | | | | Bemue-Congo | | | | | | | | | | | | | |
| | | | | | | | | Bantoid | | | | | | | | | | | | |
| | | | | | | | | | Southern Bantoid | | | | | | | | | | | |
| | | | | | | | | Northen Bantoid | | | Narrow Bantu Central | | | | | | | | | |
| cluster | Afro-Asiatic | Nilo-Saharan | Mande | Volta-Congo North | Atlantic Congo | Kwa | Others | Dakoid | Ekoid | Tivoid | E | F | G | J | K | L | M | N | P | S |
| cl1 | 9 | 6 | 0 | 3 | 0 | 0 | 1 | 0 | 0 | 0 | 1 | 0 | 0 | 1 | 0 | 1 | 1 | 1 | 0 | 2 |
| cl2a | 1 | 0 | 0 | 0 | 0 | 0 | 0 | 0 | 0 | 0 | 0 | 3 | 6 | 0 | 0 | 0 | 0 | 3 | 3 | 2 |
| cl2b | 1 | 0 | 0 | 0 | 0 | 0 | 0 | 0 | 0 | 0 | 0 | 0 | 0 | 0 | 0 | 0 | 0 | 0 | 0 | 21 |
| cl3 | 12 | 1 | 0 | 1 | 0 | 0 | 0 | 0 | 0 | 0 | 0 | 0 | 0 | 0 | 0 | 0 | 0 | 0 | 0 | 0 |
| cl4 | 17 | 0 | 0 | 0 | 0 | 0 | 0 | 0 | 0 | 0 | 0 | 0 | 0 | 0 | 0 | 0 | 0 | 0 | 0 | 0 |
| cl5a | 4 | 2 | 0 | 9 | 0 | 1 | 7 | 1 | 1 | 1 | 0 | 0 | 0 | 0 | 0 | 0 | 0 | 0 | 0 | 0 |
| cl5b | 0 | 0 | 5 | 9 | 4 | 0 | 1 | 0 | 0 | 0 | 0 | 0 | 0 | 0 | 0 | 0 | 0 | 0 | 0 | 0 |
| cl5c | 0 | 0 | 3 | 0 | 4 | 0 | 0 | 0 | 0 | 0 | 0 | 0 | 0 | 0 | 0 | 0 | 0 | 0 | 0 | 0 |
| cl6 | 0 | 6 | 0 | 0 | 0 | 0 | 0 | 0 | 0 | 0 | 0 | 1 | 1 | 16 | 0 | 0 | 0 | 0 | 0 | 0 |
| Admixed | 0 | 2 | 1 | 0 | 1 | 0 | 0 | 0 | 0 | 0 | 1 | 0 | 1 | 0 | 1 | 0 | 1 | 0 | 0 | 2 |

**Table S5.** Goodness-of-fit tests of the four models simulated in the study: models with a constant ancestral population size (Cst.) or with a bottleneck before the expansion (Bott.). The goodness-of-fit statistics measure the mean distance between the observed summary statistics and the 5,000 simulated statistics that have been accepted ($D_{prior}$) or the statistics simulated based on parameters sampled from the posterior distribution ($D_{post}$).

i) Prior predictive checks

| | | cl5c excluded | cl5c included |
|---|---|---|---|
| Cst. | *p*-value | 0.03 | 0.01 |
| | *D*prior | 13.26 | 18.9 |
| Bott. | *p*-value | 0.01 | 0 |
| | *D*prior | 13.61 | 19.16 |

ii) Posterior predictive checks

| | | cl5c excluded | cl5c included |
|---|---|---|---|
| Cst. | *p*-value | 0.03 | 0 |
| | *D*post | 7.87 | 21.84 |
| Bott. | *p*-value | 0 | 0 |
| | *Dpost* | 136.46 | 137.95 |

**Dataset S1 (separate file).** List and information of the African accessions of sorghum analyzed in this study. Available at: https://gitlab.cirad.fr/agap/sorgho/africrop_sorghum

### SI References


Abdel-Magid, A., 2003. Exploitation of food-plants in the Early and Middle Holocene Blue Nile area, Sudan and neighbouring areas. Complutum, ISSN 1131-6993, Nº 14, 2003, pags. 345-372 14, 345–372.

Abdel-Magid, A., 1989. Plant domestication in the Middle Nile Basin: an archaeoethnobotanical case study. BAR Publishing. https://doi.org/10.30861/9780860546641

Alexander, D.H., Novembre, J., Lange, K., 2009. Fast model-based estimation of ancestry in unrelated individuals. Genome Res. 19, 1655–1664. https://doi.org/10.1101/gr.094052.109

Barron, A., Fuller, D.Q., Stevens, C., Champion, L., Winchell, F., Denham, T., 2020. Snapshots in time: MicroCT scanning of pottery sherds determines early domestication of sorghum (*Sorghum bicolor*) in East Africa. Journal of Archaeological Science 123, 105259. https://doi.org/10.1016/j.jas.2020.105259

Beck, H.E., Zimmermann, N.E., McVicar, T.R., Vergopolan, N., Berg, A., Wood, E.F., 2018. Present and future Köppen-Geiger climate classification maps at 1-km resolution. Sci Data 5, 180214. https://doi.org/10.1038/sdata.2018.214

Beldados, A., Manzo, A., Murphy, C., Stevens, C.J., Fuller, D.Q., 2018. Evidence of sorghum cultivation and possible pearl millet in the second millennium BC at Kassala, Eastern Sudan, in: Mercuri, A.M., D’Andrea, A.C., Fornaciari, R., Höhn, A. (Eds.), Plants and People in the African Past: Progress in African Archaeobotany. Springer International Publishing, Cham, pp. 503–528. https://doi.org/10.1007/978-3-319-89839-1_22

Beldados, A., Ruiz-Giralt, A., 2023. Burning questions: experiments on the effects of charring on domestic and wild sorghum. Journal of Archaeological Science: Reports 51, 104170. https://doi.org/10.1016/j.jasrep.2023.104170

Boardman, S., 2000. Archaeobotany, in: Archaeology at Aksum, Ethiopia. The British Institute in Eastern Africa and The Society of Antiquaries London, London, pp. 1993-1997 363–470.

Boardman, S., 1999. The agricultural foundation of the Aksumite Empire, Ethiopia, in: van der Veen, M. (Ed.), The Exploitation of Plant Resources in Ancient Africa. Springer US, Boston, MA, pp. 137–147. https://doi.org/10.1007/978-1-4757-6730-8_12

Boivin, N., Crowther, A., Helm, R., Fuller, D.Q., 2013. East Africa and Madagascar in the Indian Ocean world. J World Prehist 26, 213–281. https://doi.org/10.1007/s10963-013-9067-4

Brass, M., Fuller, D.Q., MacDonald, K., Stevens, C., Adam, A., Kozieradzka-Ogunmakin, I., Abdallah, R., Alawad, O., Abdalla, A., Gregory, I.V., Wellings, J., Hassan, F., Abdelrahman, A., 2019. New findings on the significance of Jebel Moya in the Eastern Sahel. Azania: Archaeological Research in Africa 54, 425–444. https://doi.org/10.1080/0067270X.2019.1691845

Champion, L., Fuller, D., 2018. Archaeobotanical remains, in: Two Thousand Years in Dendi, Northern Benin: Archaeology, History and Memory. Brill, Leiden, pp. 216–233.

Costantini, L., Fattovich, R., Pardini, E., Piperno, M., 1982. Preliminary report of archaeological investigations at the site of Teglinos (Kassala). Nyame Akuma 21, 30–33.

Costantini, L., Fattovich, R., Piperno, M., Sadr, K., 1983. Gash delta archaeological project: 1982 field season. Nyame Akuma 23, 17–19.

Currat, M., Ray, N., Excoffier, L., 2004. splatche: a program to simulate genetic diversity taking into account environmental heterogeneity. Molecular Ecology Notes 4, 139–142. https://doi.org/10.1046/j.1471-8286.2003.00582.x

Denbow, J.R., Wilmsen, E.N., 1986. Advent and course of ppastoralism in the Kalahari. Science 234, 1509–1515. https://doi.org/10.1126/science.234.4783.1509

Deu, M., Hamon, P., Dufour, P., D’Hont, A., Lanaud, C., Chantereau, J., 1995. Mitochondrial DNA diversity in wild and cultivated sorghum. Genome 38, 635–645. https://doi.org/10.1139/g95-081

Deu, M., Rattunde, F., Chantereau, J., 2006. A global view of genetic diversity in cultivated sorghums using a core collection. Genome 49, 168–180. https://doi.org/10.1139/g05-092

Ehret, C., 1979. On the antiquity of agriculture in Ethiopia. The Journal of African History 20, 161–177. https://doi.org/10.1017/S002185370001700X

Fitak, R.R., 2021. OptM: estimating the optimal number of migration edges on population trees using Treemix. Biology Methods and Protocols 6, bpab017. https://doi.org/10.1093/biomethods/bpab017
Fuller, D., 2004. The Central Amri to Kirbekan survey. a preliminary report on excavations and survey 2003-04.
Fuller, D.Q., 2014. Agricultural innovation and state collapse in Meroitic Nubia: the impact of the savannah package. Left Coast Press, Walnut Creek, pp. 165–178.
Fuller, D.Q., 2004. Early Kushite agriculture: archaeobotanical evidence from Kawa. Sudan Nubia 8, 70–74.
Fuller, D.Q., Stevens, C.J., 2018. Sorghum domestication and diversification: a current archaeobotanical perspective, in: Mercuri, A.M., D’Andrea, A.C., Fornaciari, R., Höhn, A. (Eds.), Plants and People in the African Past: Progress in African Archaeobotany. Springer International Publishing, Cham, pp. 427–452. https://doi.org/10.1007/978-3-319-89839-1_19
Giblin, J.D., Fuller, D.Q., 2011. First and second millennium a.d. agriculture in Rwanda: archaeobotanical finds and radiocarbon dates from seven sites. Veget Hist Archaeobot 20, 253–265. https://doi.org/10.1007/s00334-011-0288-0
Gilabert, A., Burgarella, C., Calatayud, C., Berger, A., Rami, J.-F., Pot, D., Deu, M., 2023. Shedding light on the evolutionary history of wild and cultivated african sorghum: the guinea margaritiferum case, in: Sorghum in the 21st Century: Resiliency and Sustainability in the Face of Climate Change. Book of Abstracts. Presented at the 2023 Sorghum in the 21st Century Global Sorghum Conference, CIRAD, Montpellier, France. https://doi.org/10/609321.pdf
Hanisch, E., 1981. Schroda, a Zhizo site in the Northern Transvaal., in: Guide to Archaeological Sites in the Northern and Eastern Transvaal.
Lewis, M., 2009. Ethnologue: languages of the World, 16th edition. SIL International.
Mace, E.S., Tai, S., Gilding, E.K., Li, Y., Prentis, P.J., Bian, L., Campbell, B.C., Hu, W., Innes, D.J., Han, X., Cruickshank, A., Dai, C., Frère, C., Zhang, H., Hunt, C.H., Wang, X., Shatte, T., Wang, M., Su, Z., Li, J., Lin, X., Godwin, I.D., Jordan, D.R., Wang, J., 2013. Whole-genome sequencing reveals untapped genetic potential in Africa’s indigenous cereal crop sorghum. Nat Commun 4, 2320. https://doi.org/10.1038/ncomms3320
Madella, M., García-Granero, J.J., Out, W.A., Ryan, P., Usai, D., 2014. Microbotanical evidence of domestic cereals in Africa 7000 years ago. PLOS ONE 9, e110177. https://doi.org/10.1371/journal.pone.0110177
Maggs, T., Ward, V., 1984. Early Iron Age sites in the Muden area of Natal. Southern African Humanities 26, 105–40.
Magnavita, C., 2002. Recent archaeological finds of domesticated *Sorghum bicolor* in the Lake Chad region. Nyame akuma 14–20.
Mcintosh, S.K., 1995. Excavations at Jenne-jeno, Hambarketolo and Kaniana: the 1981 season, University of California Monographs in Anthropology. ed.
Mitchell, P., 2002. The archaeology of Southern Africa. Cambridge University Press.
Morris, G.P., Ramu, P., Deshpande, S.P., Hash, C.T., Shah, T., Upadhyaya, H.D., Riera-Lizarazu, O., Brown, P.J., Acharya, C.B., Mitchell, S.E., Harriman, J., Glaubitz, J.C., Buckler, E.S., Kresovich, S., 2013. Population genomic and genome-wide association studies of agroclimatic traits in sorghum. Proceedings of the National Academy of Sciences 110, 453–458. https://doi.org/10.1073/pnas.1215985110
Pickrell, J.K., Pritchard, J.K., 2012. Inference of population splits and mixtures from genome-wide allele frequency data. PLOS Genetics 8, e1002967. https://doi.org/10.1371/journal.pgen.1002967
Robertson, J.H., 1991. Origin and development of the Early Iron Age in south central Africa. Cincinnati, Ohio.
Ruiz-Giralt, A., Nixon-Darcus, L., D’Andrea, A.C., Meresa, Y., Biagetti, S., Lancelotti, C., 2023. On the verge of domestication: early use of C4 plants in the Horn of Africa. Proceedings of the National Academy of Sciences 120, e2300166120. https://doi.org/10.1073/pnas.2300166120
Sagnard, F., Deu, M., Dembélé, D., Leblois, R., Touré, L., Diakité, M., Calatayud, C., Vaksmann, M., Bouchet, S., Mallé, Y., Togola, S., Traoré, P.C.S., 2011. Genetic diversity, structure, gene flow and evolutionary relationships within the *Sorghum bicolor* wild–weedy–crop complex in a western African region. Theor Appl Genet 123, 1231–1246. https://doi.org/10.1007/s00122-011-1662-0

Sinclair, P.J., Morais, J.M., Adampwicz, L., Duarte, R.T., 1993. A perspective on archaeological research in Mozambique, in: The Archaeology of Africa: Food, Metals and Towns. Routledge, London, pp. 409–431.
Smith, W., 2003. Archaeobotanical investigations of agriculture at late Antique Kom El-Nana (Tell El-Amarna). Egypt Exploration Society, London.
Stemler, A., 1990. A scanning electron microscopic analysis of plant impressions in pottery from the sites of Kadero, El Zakiab, Um Direiwa and El Kadada. A Scanning Electron Microscopic Analysis of Plant Impressions in Pottery from the Sites of Kadero, El Zakiab, Um Direiwa and El Kadada 4, 87–105.
Winchell, F., Stevens, C.J., Murphy, C., Champion, L., Fuller, D.Q., 2017. Evidence for sorghum domestication in fourth millennium BC Eastern Sudan: spikelet morphology from ceramic impressions of the Butana group. Curr. Anthropol. 58, 673–683. https://doi.org/10.1086/693898
Zach, B., Klee, M., 2003. Four thousand years of plant exploitation in the Chad Basin of NE Nigeria II: discussion on the morphology of caryopses of domesticated Pennisetum and complete catalogue of the fruits and seeds of Kursakata. Veget Hist Archaeobot 12, 187–204. https://doi.org/10.1007/s00334-003-0016-5
